# Multimodal single-cell analysis of non-random heteroplasmy distribution in human retinal mitochondrial disease

**DOI:** 10.1101/2022.06.20.496449

**Authors:** Nathaniel K. Mullin, Andrew P. Voigt, Miles J. Flamme-Wiese, Xiuying Liu, Megan J. Riker, Katayoun Varzavand, Edwin M. Stone, Budd A. Tucker, Robert F. Mullins

**Author notes:** Address correspondence to: Robert F. Mullins 4130 MERF 375 Newton Rd Iowa City, IA 52242 USA.

## Abstract

Variants within the high copy number mitochondrial genome (mtDNA) can disrupt organelle function and lead to severe multi-system disease. The wide range of manifestations observed in mitochondrial disease patients results from varying fractions of abnormal mtDNA molecules in different cells and tissues, a phenomenon termed heteroplasmy. However, the landscape of heteroplasmy across cell types within tissues and its influence on phenotype expression in affected patients remains largely unexplored. Here, we identify non-random distribution of a pathogenic mtDNA variant across a complex tissue using single-cell RNA sequencing, mitochondrial single-cell ATAC sequencing, and multimodal single-cell sequencing. We profile the transcriptome, chromatin accessibility state, and heteroplasmy in cells from the eyes of a patient with mitochondrial encephalopathy, lactic acidosis, and stroke-like episodes (MELAS) and healthy control donors. Utilizing the retina as a model for complex multi-lineage tissues, we found that the proportion of the pathogenic m.3243A>G allele was neither evenly nor randomly distributed across diverse cell types. All neuroectoderm-derived neural cells exhibited a high percentage of the mutant variant. However, a subset of mesoderm-derived lineage, namely the vasculature of the choroid, was near homoplasmic for the wildtype allele. Gene expression and chromatin accessibility profiles of cell types with high and low proportions of m.3243A>G implicate mTOR signaling in the cellular response to heteroplasmy. We further found by multimodal single-cell sequencing of retinal pigment epithelial cells that a high proportion of the pathogenic mtDNA variant was associated with transcriptionally and morphologically abnormal cells. Together, these findings show the non-random nature of mitochondrial variant partitioning in human mitochondrial disease and underscore its implications for mitochondrial disease pathogenesis and treatment.

**GRAPHICAL ABSTRACT:** 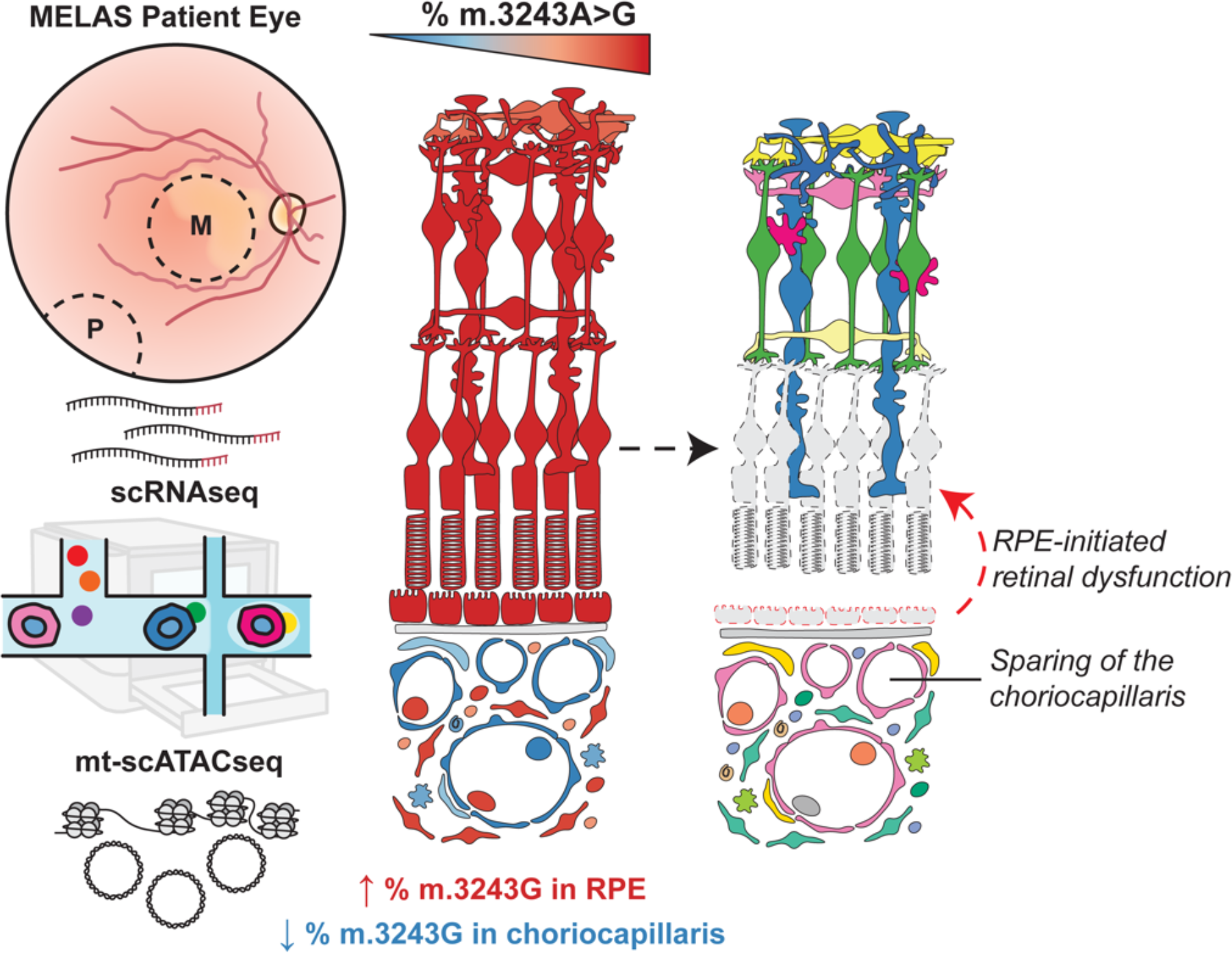

## INTRODUCTION

Several single nucleotide variants on the mitochondrial genome are known to cause retinal dystrophy and multi-system disease (1). A notable feature of these variants is that they exhibit a high degree of phenotypic heterogeneity (2). This heterogeneity manifests in terms of timing of disease onset, organ systems involved, and severity of tissue dysfunction (3, 4). The heterogenous nature of genetic mitochondrial disease is likely due, at least in part, to the fact that the mitochondrial genome exists at a high copy number in relation to the nuclear genome (5). Since every cell possesses hundreds to thousands of mitochondrial genomes (mtDNA) (6), individual cells may contain a mixture of mutant and wildtype alleles, a phenomenon known as heteroplasmy (7). Mutant mitochondrial alleles are known to be unevenly distributed among organs (8, 9) and cells within the same organ (10, 11). How the distribution of mutant and wildtype mitochondrial molecules is established during organogenesis and maintained throughout life, as well as influence of this distribution on the resulting disease phenotype, are areas of active interest in the study of mitochondrial disease (3, 5, 12, 13).

Here, we focus on the most common pathogenic mitochondrial variant: m.3243A>G, a transition mutation found in the mitochondrial leucine tRNA gene *MT-TL1* (14). m.3243A>G is thought to disrupt electron transport chain function through mistranslation of the thirteen protein-coding genes found on the mtDNA (15–20).

Depending on the age of onset and organ system involvement, patients with this variant are typically diagnosed with one of three syndromic conditions: maternally inherited diabetes and deafness (MIDD, OMIM 520000), mitochondrial encephalopathy, lactic acidosis, and stroke-like episodes (MELAS, OMIM 540000) or Leigh syndrome (OMIM 256000) (20, 21). Symptoms can include vision loss (22–24), sensorineural hearing loss (25), diabetes mellitus, gastrointestinal disturbances, myopathy, stroke-like episodes (20), and premature death (26). Notably, the affected organs in these syndromes arise from all three primordial germ layers. In addition to causing dysfunction in a broad range of organ systems, the phenotypic expression of m.3243A>G is also highly variable, even within affected families (27). The heterogenous pattern of organ involvement between patients with the m.3243G variant has led to speculation that disease may be explained by the random partitioning of the pathogenic allele during early development, such that affected organs are simply those that randomly contain a higher proportion of m.3243G. However, emerging evidence has called some of the assumptions underpinning this model of m.3243A>G pathogenesis into question. Global mutant allele burden is not sufficient to explain the heterogeneity observed in m.3243A>G associated disease phenotype (28, 29), implying a role for variable intra-tissue heteroplasmy. Recent work in circulating blood suggests that allelic segregation within tissues can be non-random (11). The concept of controlled mitochondrial DNA distribution builds on previous observations that mutant allele proportion followed predictable patterns between tissues (21, 30) and that such patterns may even be heritable, implying the involvement of nuclear genetic factors in governing the segregation or propagation of mutant alleles (3, 29).

In the current study, our goal was to investigate the relationship between the distribution of m.3243A>G across the retina and the clinical retinal phenotype in MELAS. We sought to test whether m.3243G was evenly distributed among the cell types of the human retina and to examine the effect of pathogenic variant dosing in specific ocular cell types on cellular phenotype. To accomplish this, we utilized primary post-mortem human ocular tissue from patients diagnosed with m.3243A>G associated MELAS. From this tissue, we collected neural retina, retinal pigment epithelium (RPE), and choroid (31). The retina, RPE, and choroid are an ideal system for the study of the cell type-specific dynamics that govern heteroplasmy and cellular response to a mitochondrial variant in a complex tissue as derivatives of neuroectoderm and mesoderm are in close physical and functional association. We provide evidence that the ratio m.3243G to m.3243A is not randomly distributed across the cell types of the neural retina and choroid. We identify dysregulation of transcriptional targets downstream of mTOR signaling that may act as potential mediators of heteroplasmy modulation *in vivo*.

Understanding the process of heteroplasmy control by cells *in vivo* could enable heteroplasmy modulation as a therapeutic approach. Together, combining these intra-organ heteroplasmy data with prior knowledge of retinal metabolism and clinical manifestations of mitochondrial disease, we gain an increased understanding of m.3243A>G pathogenesis in the eye.

## RESULTS

### Clinical ophthalmologic history of m.3243A>G disease

Although m.3243A>G is known to be sufficient to cause disease (14, 32, 33), the sequence of events that connect the presence of the mtDNA mutation in cells to the widespread retinal atrophy is not fully understood (3, 29). Fundus examination of the proband revealed macular atrophy typical of the m.3243A>G variant (**Figure 1A, B**) (22). Optical coherence tomography (OCT) of the retina showed outer retinal tubulations (ORTs) (**Figure 1C**), a clinical finding known to be associated with m.3243A>G associated disease and with RPE/choroid dysfunction leading to retinal dystrophy more broadly (34–36). The pedigree structure was compatible with mitochondrial inheritance and the proband had been previously confirmed to carry the m.3243A>G variant by sequencing and digital PCR (**Figure S1A**). This patient was shown previously to harbor a relatively high global burden of the m.3243A>G variant in blood cells and skin fibroblasts compared to patients with less severe disease manifestations (30). Intra-tissue heteroplasmy distribution has not previously been explored in the eye by ourselves or others.

**Figure 1.**
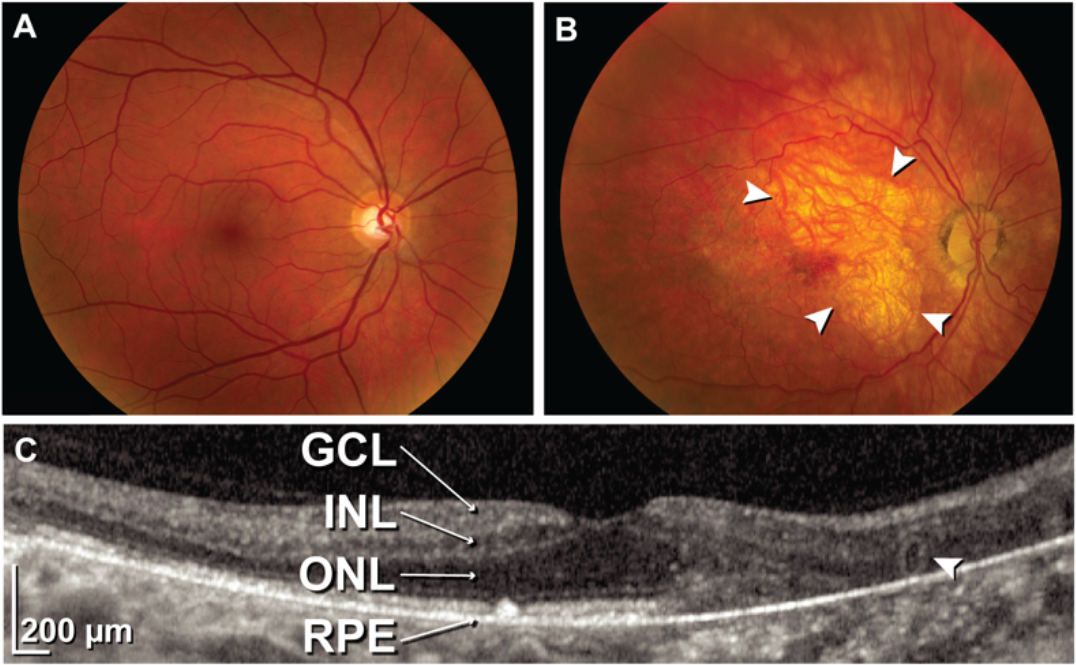
Macular retinopathy caused by the m.3243A>G variant. **(A)** Color fundus photograph of the right eye of a healthy control patient with 20/20 acuity. **(B)** Color fundus photograph of the proband. Macular atrophy of the RPE and choriocapillaris typical of MELAS is appreciable (white arrowheads). **(C)** Optical coherence tomography (OCT) of the proband demonstrates outer retinal tubulations (white arrowhead) indicative of migration of cone photoreceptor cells following RPE cell death (GCL; ganglion cell layer, INL; inner nuclear layer, ONL; outer nuclear layer, RPE; retinal pigment epithelium/Bruch’s complex).

### Single cell profiling of MELAS retina and choroid

To better understand how the proportion of m.3243G varies within tissues and how such variability affects cellular dysfunction, we utilized a microfluidic-based single-cell sequencing approach to profile the transcriptome, nuclear chromatin accessibility, and heteroplasmy of single cells isolated from the neural retina and underlying RPE and choroid. Cells were isolated from the right eye (OD) of a patient with clinically and genetically diagnosed MELAS. As m.3243A>G is known to cause region-specific retinal atrophy (22), samples were recovered from both the macular (M) and peripheral (P) retina, as well as from that of an age and sex-matched control individual. The neural retina and underlying RPE & choroid were each dissected, dissociated, and profiled with scRNAseq and mt-scATACseq (37) (**Figure 2A**). Cells profiled by single-cell RNA sequencing (scRNAseq) were projected in a low dimensional space based on gene expression (**Figure 2B, C**), and resulting clusters were manually annotated using previously described marker genes for human ocular cell types (38, 39) (**Figure 2D, E**). Due to the sparseness inherent in the single cell assay of transposase-accessible chromatin by sequencing (scATAC-seq) data, we annotated mt-scATAC-seq clusters via unsupervised label transfer from the corresponding scRNA-seq datasets (**Figure 2F, G**) using previously described methods (40). Clusters were examined to ensure proper integration of both control and proband samples and between cells taken from the macula or periphery (**Figure S1E-H**). The number of cells analyzed in each sample using either modality is shown in **Table 1**.

**Figure 2.**
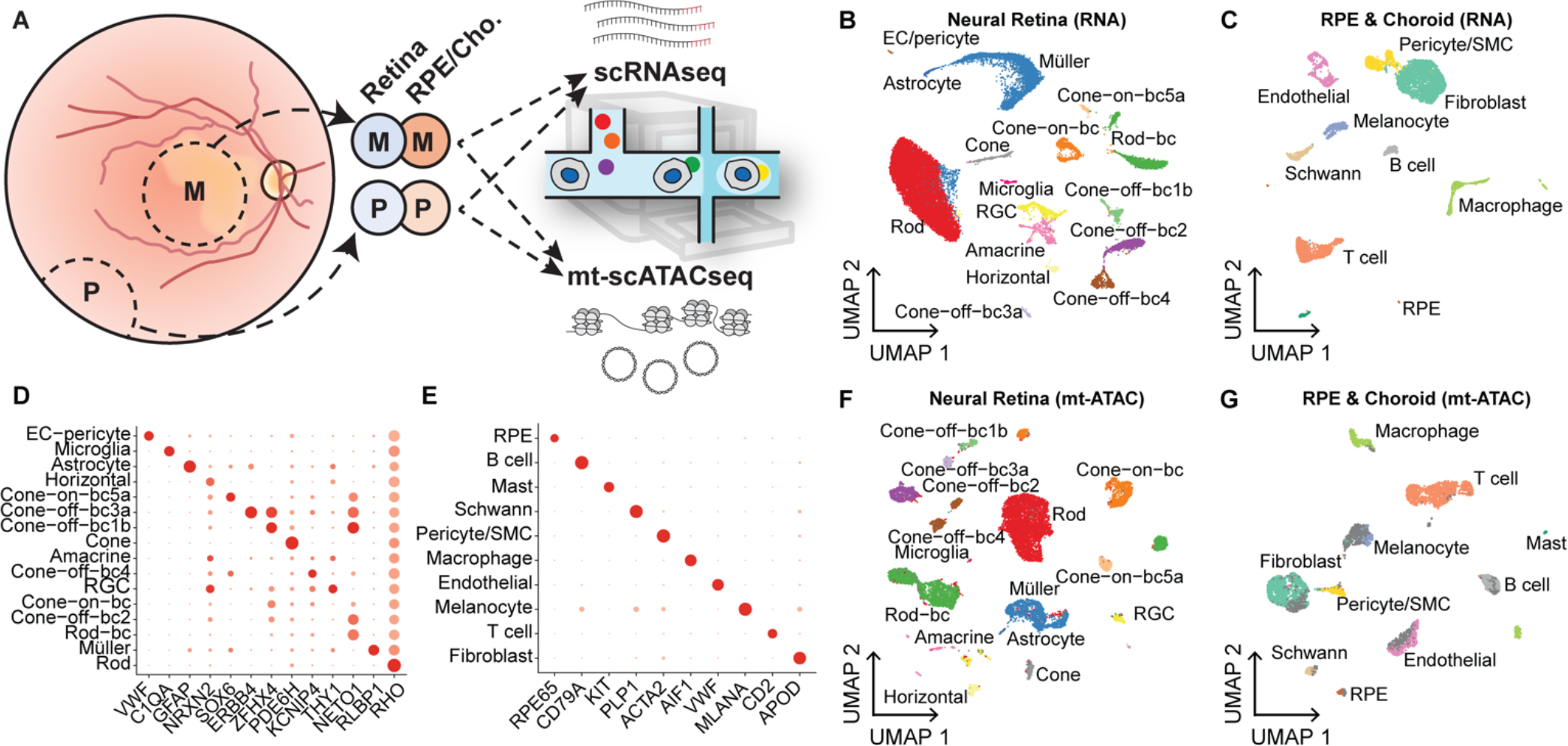
Transcriptome and chromatin accessibility profiling in single cells isolated from MELAS and control eyes. **(A)** Experimental schematic of scRNAseq and mt-scATACseq studies. Macular (M) and peripheral (P) punches were dissected into retina and RPE/choroid samples. These samples were dissociated, split, and subjected to scRNAseq and mt-scATACseq. **(B)** Two-dimensional UMAP embedding of neural retinal cells based on gene expression (scRNAseq) data from the proband and control donor. **(C)** Two-dimensional UMAP embedding of RPE, and choroidal cells based on scRNAseq data from the proband and control donor. **(D)** Dot plot indicating the magnitude of expression (shade of red) and proportion of cells (size of dot) expressing known marker genes in the neural retinal clusters annotated in (B). **(E)** Dot plot of curated marker genes in the RPE/choroid clusters annotated in (C). **(F)** Two-dimensional UMAP embedding of neural retinal cells based on chromatin accessibility (mt-scATACseq) data from the proband and control donor. **(G)** Two-dimensional UMAP embedding of RPE and choroidal cells based on mt-scATACseq data from the proband and control donor. Cells with label transfer prediction score less than 0.6 are shaded in dark gray. (EC; endothelial cell, BC; bipolar cell, RGC; retinal ganglion cell, SMC; smooth muscle cell, RPE; retinal pigment epithelium).

**Table 1.**
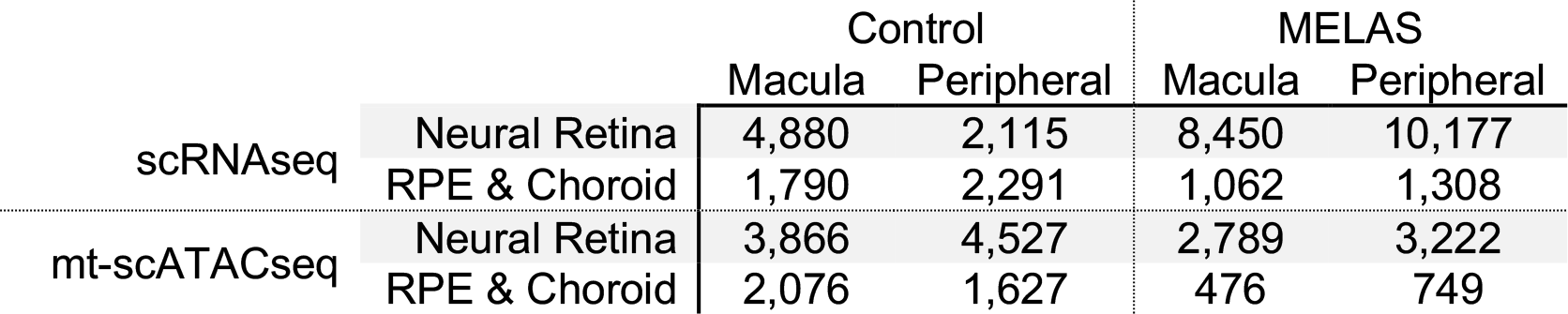
Cell counts from MELAS and control individuals profiled by scRNAseq and mt-scATACseq.

|  |  | Control |  | MELAS |  |
| --- | --- | --- | --- | --- | --- |
|  |  | Macula | Peripheral | Macula | Peripheral |
| scRNAseq | Neural Retina | 4,880 | 2,115 | 8,450 | 10,177 |
|  | RPE & Choroid | 1,790 | 2,291 | 1,062 | 1,308 |
| mt-scATACseq | Neural Retina | 3,866 | 4,527 | 2,789 | 3,222 |
|  | RPE & Choroid | 2,076 | 1,627 | 476 | 749 |

No major region-specific changes in gene expression were observed and cells from these regions were computationally pooled for subsequent analyses. Since scRNA-seq and mt-scATAC-seq datasets were derived from the same initial samples, a high correlation between cell type proportion was observed (**Figure S1B**) lending confidence to the unsupervised label transfer approach. Confidence scores for predicted cell type classification were generated (**Figure S1C, D**), and only cells with a prediction score greater than 0.6 were used in downstream analysis. Cells not reaching this prediction score are colored in dark grey in **Figure 2F and G**.

### Non-random m.3243G partitioning in the retina and choroid

Since the partitioning of the m.3243A>G variant within tissues would likely affect the resulting clinical phenotype, we asked whether heteroplasmy of the m.3243A>G variant differed among cells isolated from the neural retina and RPE/choroid (**Figure 3A**). mt-scATACseq data were used, and only cells with sufficient coverage of the m.3243 locus were included in the analysis (see **Methods**). In order to confirm that neither coverage of ATAC reads nor depth of mtDNA sequencing confounded heteroplasmy determination in single cells, these variables were plotted against m.3243G proportion for each cell included in **Figure 3** (**Figure S2).** No association between heteroplasmy calls and locus (**Figure S2A-B**) or mtDNA (**Figure S2C-D**) coverage was observed. The per-cell proportion of pathogenic m.3243G allele was overlayed onto cells projected in a low dimensional space (**Figure 3B, C**). Major cell types of the neural retina (photoreceptor cells, interneurons, and glia) were near homoplasmic for the pathogenic m.3243G allele (**Figure 3B**), all having median m.3243G proportions of 1. This finding was corroborated by bulk dPCR analysis of the neural retina (**Figure S1A**). The RPE cells and fibroblasts of the RPE/choroid sample also had median m.3243G proportions of 1 (**Figure 3C**). However, endothelial and T cell populations were near homoplasmic for wildtype m.3243A, all having median m.3243G proportions beneath 0.05 (**Figure 3C**). The proportion of the pathogenic m.3243G allele within individual cells grouped by cell type is plotted in **Figure 3D** and **E**. The melanocytes and pericytes from the RPE/choroid sample were heterogenous in terms of proportion m.3243G (**Figure 3E**).

**Figure 3.**
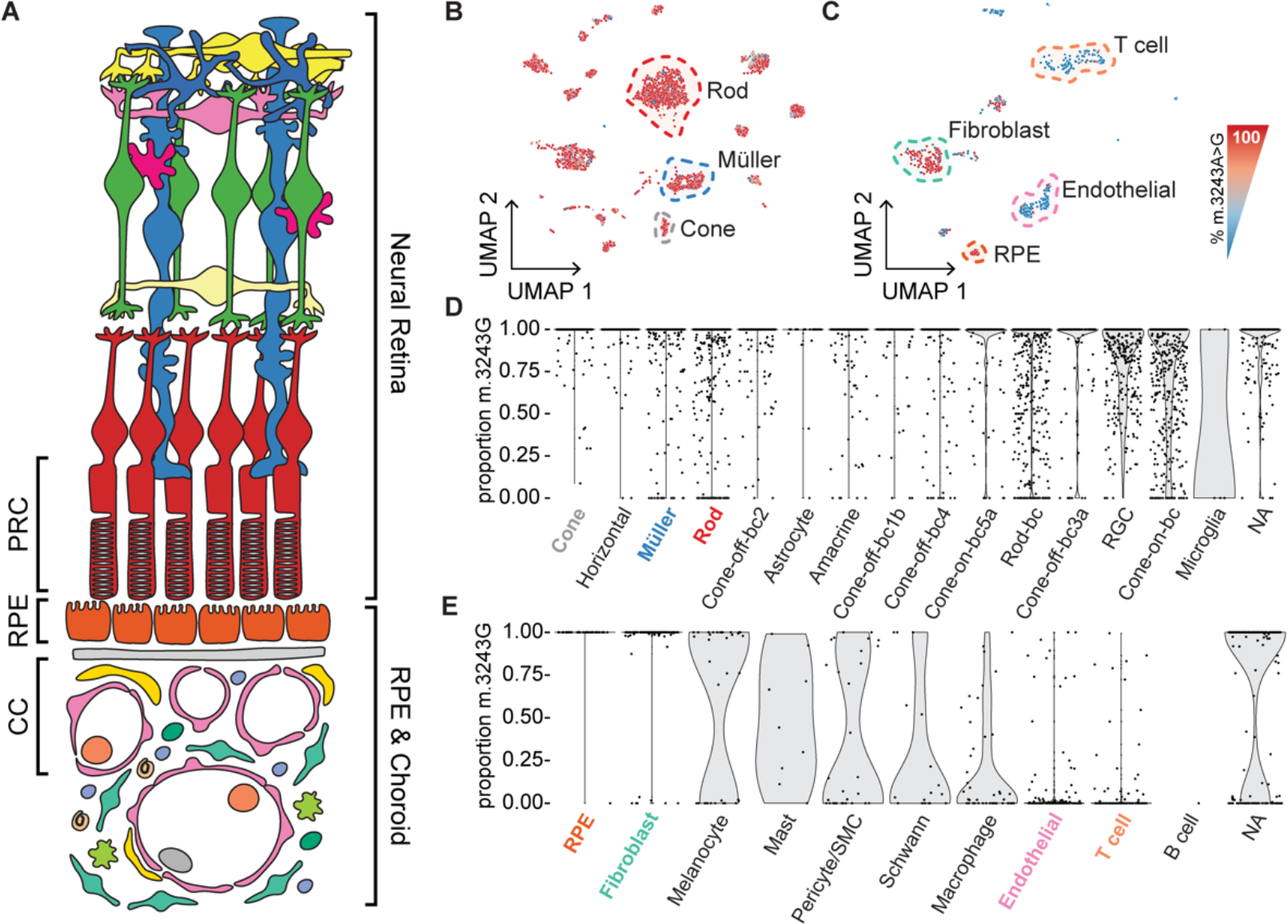
Non-random distribution of m.3243G in the retina and choroid measured by mt-scATACseq. **(A)** Schematic of the neural retina, RPE, and choroid indicating the cell classes captured by scRNAseq, mt- scATACseq, and LCM-dPCR. **(B)** Two-dimensional UMAP embeddings of cells recovered from the MELAS neural retina. Individual cells are colored based on proportion of m.3243A>G mutant allele (red indicating 100% mutant allele and blue indicating 100% wildtype allele). **(C)** Two-dimensional UMAP embeddings of cells recovered from the MELAS RPE and choroid colored based on proportion of m.3243A>G mutant allele. **(D)** Violin plot demonstrating the proportion of m.3243A>G mutant allele in individual cells isolated from the neural retina sample as measured by mt-scATACseq. Cells are grouped based on cell type cluster identity. **(E)** Violin plot demonstrating the proportion of m.3243A>G mutant allele in individual cells isolated from the RPE/choroid sample as measured by mt-scATACseq. Cells with at least 10x coverage of the m.3243 locus are included in **(D)** and **(E)**.

To validate the finding that choroidal endothelial cells have a reduced proportion of m.3243G compared to other choroidal and retinal cell types, we performed laser capture microdissection followed by digital PCR (LCM-dPCR) on independent m.3243A>G ocular samples. Photoreceptor (PRC), RPE, and choriocapillaris (CC) layers (see **Figure 4A**) were separately captured from the contralateral eye of the proband analyzed by mt-scATAC-seq (MELAS 1*) as well as from an unrelated MELAS patient known to carry the m.3243A>G variant (MELAS 2) (**Figure 4A-F**). The CC showed routinely lower m.3243A>G than the RPE and PRC layers (**Figure 4G**), consistent with the results of the mt-scATAC-seq approach (**Figure 3B-E**). Due to the highly complex and dense nature of the choroid, isolation of pure endothelial cells is not possible with LCM-dPCR, explaining the relatively elevated heteroplasmy compared to the mt-scATACseq approach.

**Figure 4.**
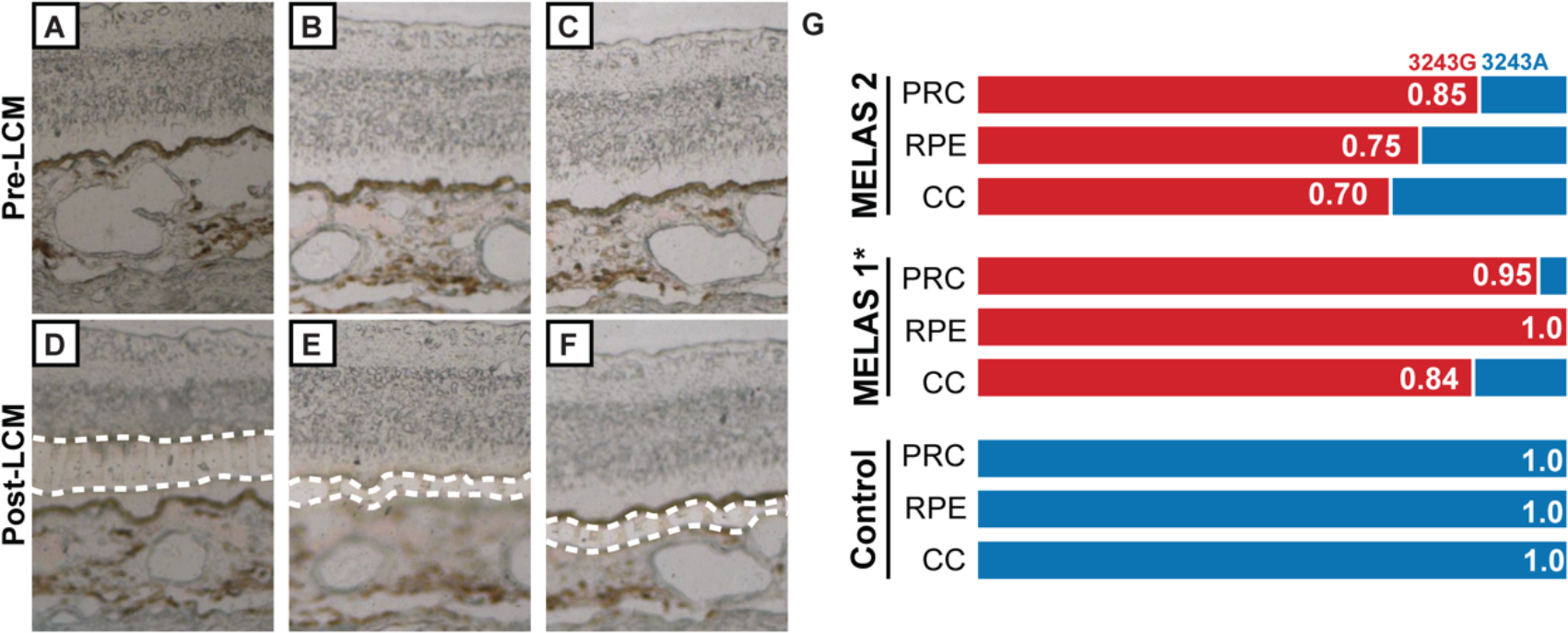
m.3243G partitioning in the retina and choroid measured by digital PCR. **(A-C)** Color photomicrographs of neural retina, RPE, and choroidal regions of fixed ocular sections. Regions of interest are shown before laser capture microdissection (*Pre-LCM*). **(D-F)** Color photomicrographs of fixed ocular sections after isolation of photoreceptors (PRC) **(D)**, RPE **(E)**, and choriocapillaris (CC) **(F)** by LCM (*Post-LCM)*. Regions isolated for subsequent digital PCR are indicated with white dotted lines. **(G)** Proportion of mutant m.3243G in each LCM-captured region as measured by digital PCR. *MELAS 1\** indicates proband eye contralateral to that used in the mt-scATAC study. *MELAS 2* is an unrelated MELAS patient.

We used transmission electron microscopy (TEM) to evaluate the effect of m.3243A>G heteroplasmy on the ultrastructural phenotype in retinal cells. We found the mitochondria in MELAS neural retina (**Figure 5A**) and RPE (**Figure 5C**) to be swollen and hypertrophied compared to those in a control sample (**Figure 5B, D**).

**Figure 5.**
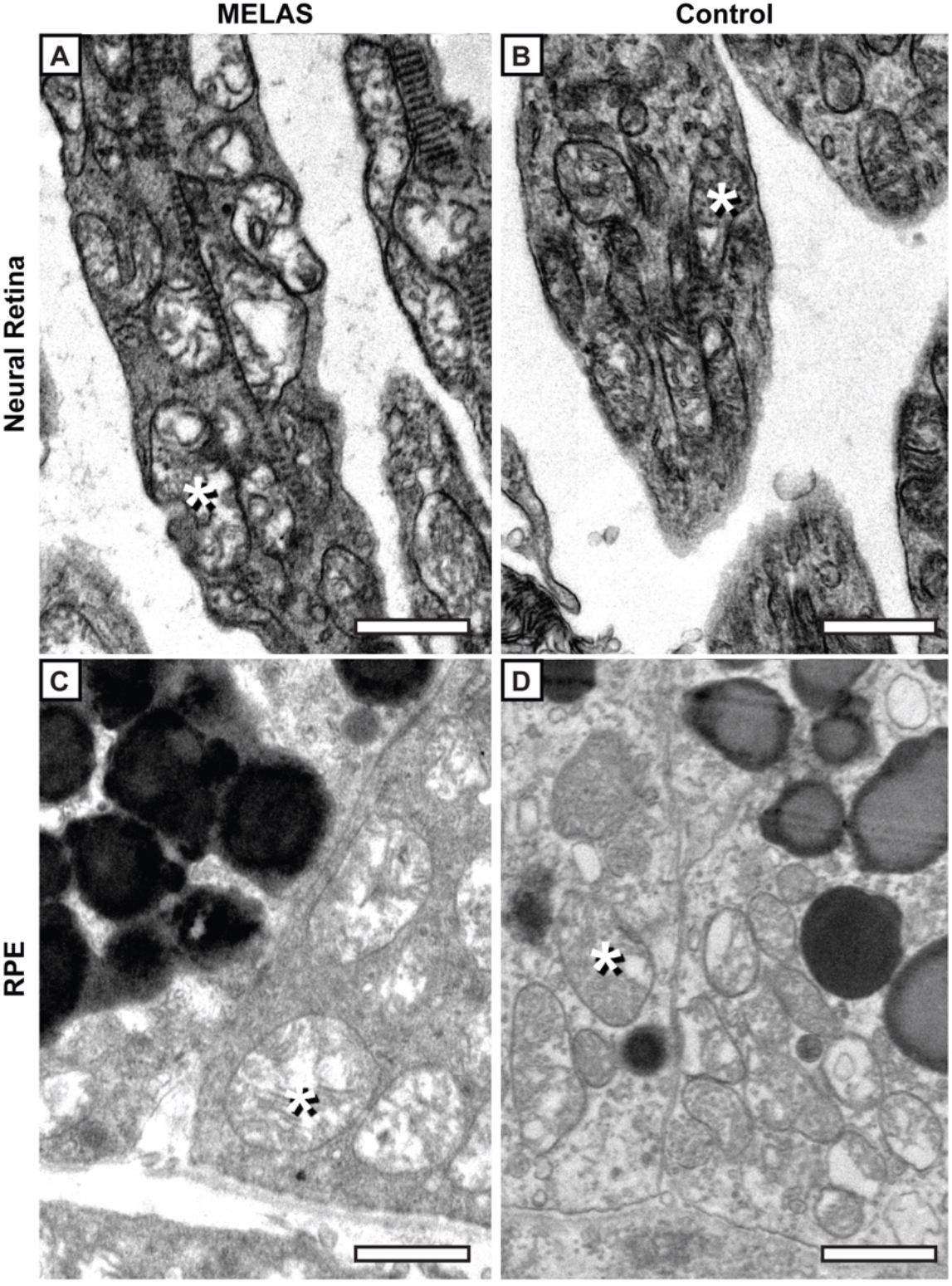
m.3243G partitioning in the retina and choroid affects mitochondria ultrastructure. **(A)** Transmission electron micrographs of photoreceptor cell inner segments from MELAS patient samples reveal abnormal mitochondrial structure compared to control tissue **(B)**. **(C)** Transmission electron micrograph of RPE cells from MELAS patient samples also show abnormal mitochondrial structure compared to control (**D**). Mitochondria are indicated with asterisks. Scale bar represents 1µm.

### Dysregulated gene expression in retinal cell types with varying m.3243A>G proportion

After determining the non-random partitioning of m.3243A>G across the cell types of the neural retina and RPE/choroid, we next asked how the presence of pathogenic m.3243G impacts gene expression in different cell types. We performed differential expression analysis of cell types from the neural retina and RPE/choroid obtained from the MELAS proband and a normal control individual. We focused on two cell types with high levels of m.3243G (i.e., cone photoreceptors and choroidal fibroblasts), and two cell types with low levels of m.3243G (i.e., T lymphocytes and choroidal endothelial cells). We observed differential gene expression in both high and low-m.3243G cell types (**Figure 6A-D**). The nuclear encoded gene *MTRNR2L1* was upregulated in the three choroidal cell types in the MELAS samples compared to controls (**Figure 6B-C,** highlighted in yellow), irrespective of cell type-specific heteroplasmy. *MTRNR2L1* encodes the peptide humanin-like 1, a peptide that is highly homologous to humanin, which is encoded by the mitochondrial gene *MT-RNR2*.

**Figure 6.**
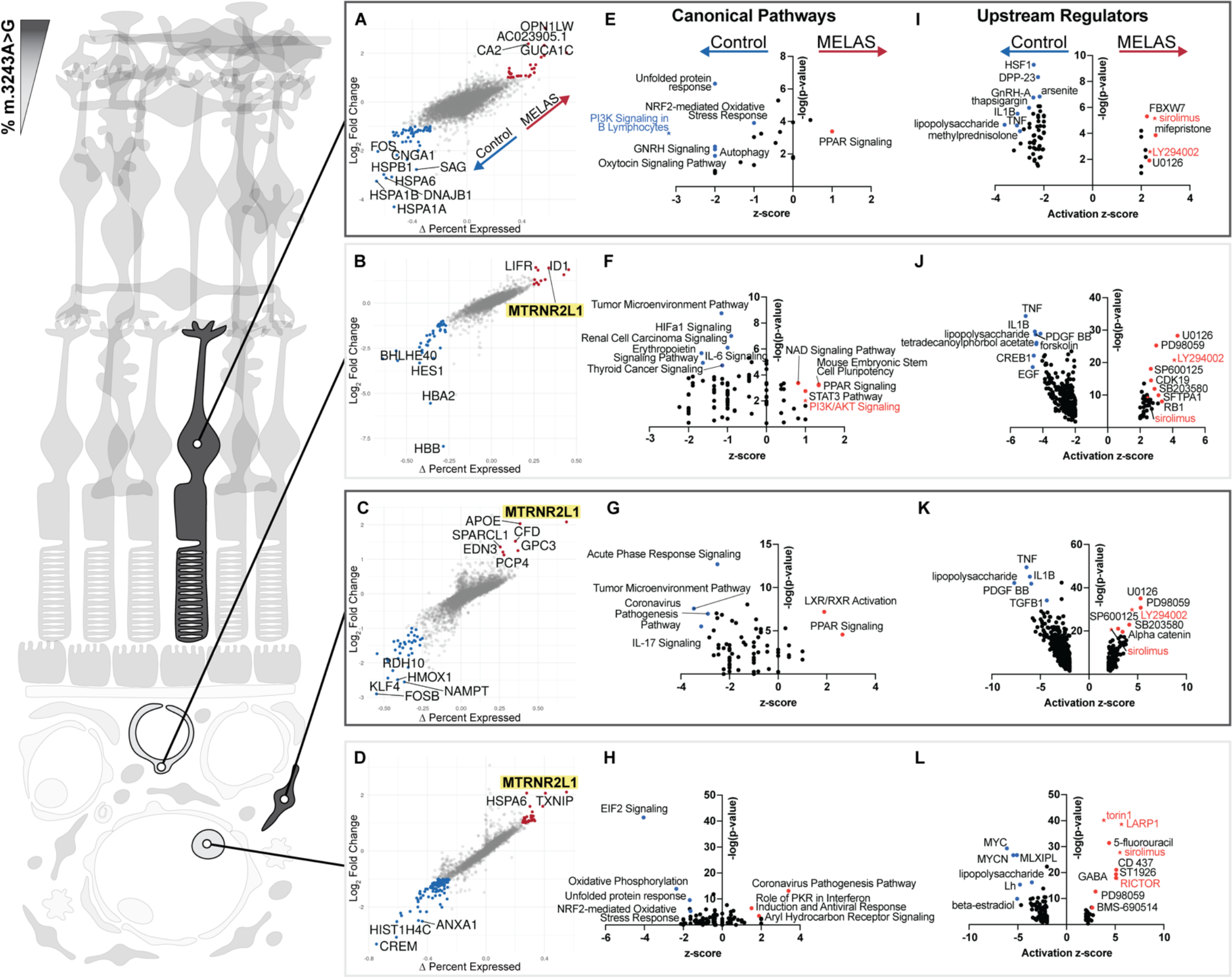
Impact of m.3243A>G heteroplasmy on retinal cell types. **(A-D)** Differentially expressed genes between MELAS and control samples in cone photoreceptors (A), choroidal endothelial cells (B), choroidal fibroblasts (C), and T lymphocytes (D). Genes expressed with over 1.5-fold change between samples and Δ percent expressed greater than 25% are colored. The most highly differentially expressed genes are labeled. Of note, *MTRNR2L1* transcript is upregulated in MELAS samples in cell types with both high (C) and low (B, D) proportions of m.3243G. **(E-H)** Canonical pathways enriched in the dysregulated genes set are plotted. Pathways involving PI3K/AKT signaling are colored in blue and red in cone photoreceptor and endothelial cell subsets (E, F). **(I-L)** Predicted upstream regulators based on the cell type-specific gene list of dysregulated genes in MELAS versus control. Plotted molecules are experimentally shown to produce gene expression changes similar to those observed in differential expression analysis. The molecules torin1, LARP1, sirolimus, RICTOR, and LY294002 are known to act on the mTOR pathway are highlighted in red in I-L.

Humanin is known to be cytoprotective against oxidative stress and has been shown to rescue RPE cells from oxidative damage *in vitro* (41).

In order to understand the functional significance of dysregulated genes in the MELAS retina and choroid, we performed pathway analysis, looking for enrichment of canonical pathway terms (**Figure 6E-H**) and for targets of known upstream regulatory factors or small molecules (**Figure 6I-L**). Pathways involved in oxidative stress response (“NRF2-mediated Oxidative Stress Response”), oxidative phosphorylation (“Oxidative Phosphorylation”), and the response to hypoxia (“HIFa1 Signaling”, “Erythropoietin Signaling Pathway”) were enriched in the control sample versus the MELAS sample in cell types with both high and low heteroplasmy (**Figure 6E-H**). The PI3K pathway was down regulated in high m.3243A>G proportion cone photoreceptors (**Figure 6E**), while being enriched in the low mutational burden choroidal endothelial cells (**Figure 6F**).

Dysregulation of PI3K signaling, along with downstream AKT and mTOR signaling, has been shown previously *in vitro* to have a role in purifying selection against mitochondria carrying the m.3243A>G allele (42). Similarly, we found that the top Upstream Regulators that recapitulate the gene expression observed in MELAS cells involved inhibition of mTOR/PI3K/AKT signaling (**Figure 6I-L**). These data indicate that the same pathway may play a role related to the control of pathogenic mtDNA alleles in the eye.

### Cell type-specific chromatin accessibility changes in MELAS

Chromatin remodeling has previously been implicated in the cellular phenotype caused by m.3243A>G (43). We sought to use our mt-scATACseq data to understand how the m.3243G variant alters transcription factor binding in retinal and choroidal cells. We used chromVAR (44) to identify transcription factor-binding motifs that were differentially enriched in MELAS versus control cells in specific choroidal cell types (**Figure 7**).

**Figure 7.**
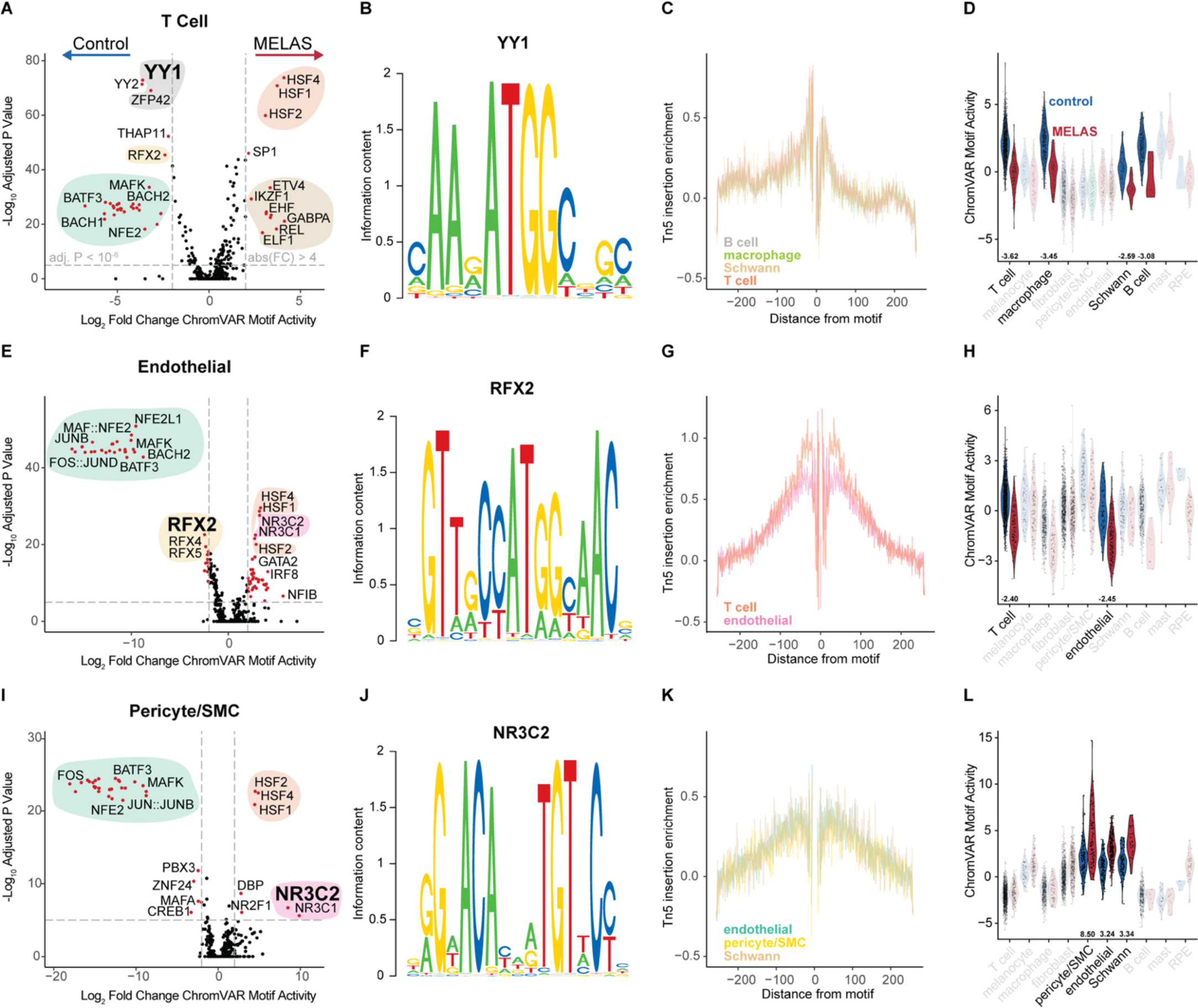
Differentially accessibility of transcription factor binding motifs in MELAS choroid. **(A)** Top motifs enriched in MELAS or control T cells. Clusters of similar motifs (as defined by JASPAR Core Vertebrate motif clusters) are shown with colored circles. The YY1 and ZFP42 motifs (Cluster 84) are de-enriched in MELAS T cells. **(B)** Motif logo of the JASPAR YY1 motif (MA0095.2). **(C)** Motif foot printing plot shows Tn5 transposase insertion enrichment in the locus flanking the YY1 binding motif. Traces for four cell types with lower YY1 motif enrichment in MELAS are shown. **(D)** YY1 binding motif enrichment score shown for all choroidal cell types. Four cell types with low m.3243A>G heteroplasmy have lower YY1 binding motif enrichment in MELAS versus control. **(E)** Enriched motifs in choroidal endothelial cells. **(F)** RFX2 motif logo, one of several RFX factor motifs enriched in control endothelial cells versus controls. **(G)** RFX2 motif footprint plot showing traces for T cells and endothelial cells. **(H)** ChromVAR activity for the RFX2 motif is lower in MELAS endothelial and T cells in MELAS sample versus control. **(I)** Enriched motifs in the pericyte and SMC population of MELAS versus control. **(J)** The NR3C2 binding motif, which is overrepresented in MELAS cells with low m.3243G abundance. **(K)** NR3C2 footprint plot indicating transcription factor-chromatin interaction in cell types of interest. **(L)** ChromVAR activity of NR3C2 is elevated in the low-heteroplasmy pericyte/SMC, endothelial, and Schwann cell populations in MELAS versus control.

Specifically, we performed differential enrichment analysis for MELAS versus control cells in the T cell (**Figure 7A**), endothelial cell (**Figure 7E**), and pericyte/SMC (**Figure 7I**) populations of the choroid. T cells and endothelial cells of the choroid exhibited near homoplasmy for the wildtype m.3243A allele, while the pericyte/SMC population displayed heteroplasmy for the m.3243A>G variant (**Figure 3E**).

We performed differential enrichment analysis of transcription factor binding motifs in MELAS versus control T cells and found YY1 to be a top hit. YY1 is a zinc-finger transcription factor known to work downstream of mTOR signaling to regulate mitochondrial function through transcription of nuclear-encoded mitochondrial genes(45). The YY1 binding motif (**Figure 7B**) and similar motifs showed decreased enrichment in T cells from the MELAS patient as compared to control (**Figure 7A**). These motifs were not differentially accessible in the homoplasmic endothelial population (**Figure 7E**). Footprint analysis of Tn5 insertion around the YY1 motif showed expected patten of reduced accessibility at the motif, indicating blocking of transposase activity by bound YY1 (**Figure 7C**). YY1 motif enrichment was lower in MELAS than control in T cell, macrophage, Schwann cell, and B cell populations (**Figure 7D**), all of which had low proportions of pathogenic m.3243G allele (**Figure 3E**).

We next performed similar analysis on the endothelial (**Figure 7E-H**) and pericyte/SMC (**Figure 7I-L**) populations. We found that T cells and endothelial cells share reduced RFX2 motif enrichment in the MELAS sample as compared to control (**Figure 7H**). Pericyte/SME, endothelial, and Schwann cell populations were all enriched for the NR3C2-binding motif in MELAS versus control cells (**Figure 7I-L)**. This motif was not enriched in T cells. NR3C2 activity has been previously linked to PI3K/ATK signaling (46), further supporting a connection between heteroplasmy modulation and signaling downstream of mTOR in the eye. Chemical inhibition of PI3K/AKT with LY294002 has been shown to reduce the proportion of m.3243G *in vitro* and to increase activity of NR3C2. Together, these analyses suggest that transcription factors acting downstream of mTOR signaling have differential activity depending on MELAS disease state and cell type.

We further identified additional motifs that were enriched in a cell type-specific manner across the retina and choroid (**Figures S3 and S4**) and found highly enriched, cell type-specific motifs in the neural retina (shown in **Figure S3A)**. One motif, MA0476.1, was enriched only in a subset of Müller cells (**Figure S3A, box and motif logo**). Results were confirmed by examining expression of transcription factor-encoding genes in the same cell type using scRNAseq data (**Figure S3B-G).** Together, these analyses confirm that motif enrichment in mt- scATACseq data represent legitimate regulation of gene expression.

### Multimodal sequencing reveals aberrant gene expression profile in MELAS RPE

RPE dysfunction has been implicated in the pathogenesis of m.3243A>G-associated vision loss (23, 47) and the outer retinal tubulations frequently observed in this condition (**Figure 1C**) are indicative of RPE cell death (48). Since the choroidal cells most responsible for photoreceptor health (endothelial cells) were homoplasmic for wildtype m.3243, we hypothesized that the RPE is the site of the initial tissue injury in the eye. To confirm the high proportion of mutant m.3243G in the RPE and investigating its impact on gene expression, we jointly profiled heteroplasmy, gene expression, and chromatin accessibility in the same single cells from choroidal and purified RPE samples of the MELAS proband and controls (**Figure 8A**). Due to a low RPE cell count in unselected RPE/choroid samples (**Figure 2B, C, F, G)**, manually purified RPE samples were included in this experiment to guarantee sufficient cell number. Cell clustering was performed using WNN-analysis (49), which combines gene expression and chromatin accessibility modalities in a weighted manner to drive and refine cluster identification (**Figure 8B**) beyond that derived from single modalities (**Figure S3A-C**). The relative modality weighting behind each cluster is shown (**Figure S3D**). While MELAS and control samples generally clustered together across most cell types (**Figure S3E**), the RPE cluster was fragmented (**Figure 8B, inset**) revealing a unique MELAS RPE sub-cluster. The RPE identity of each sub-cluster was confirmed by expression of the highly restricted marker gene *BEST1* (**Figure 8C**). Confirming prior mt-scATACseq analysis, the MELAS proband RPE was homoplasmic for mutant m.3243G (**Figure 8D, E**). We next sought to understand how this high proportion of m.3243G in the RPE related to cellular phenotype and overall retinal function.

**Figure 8.**
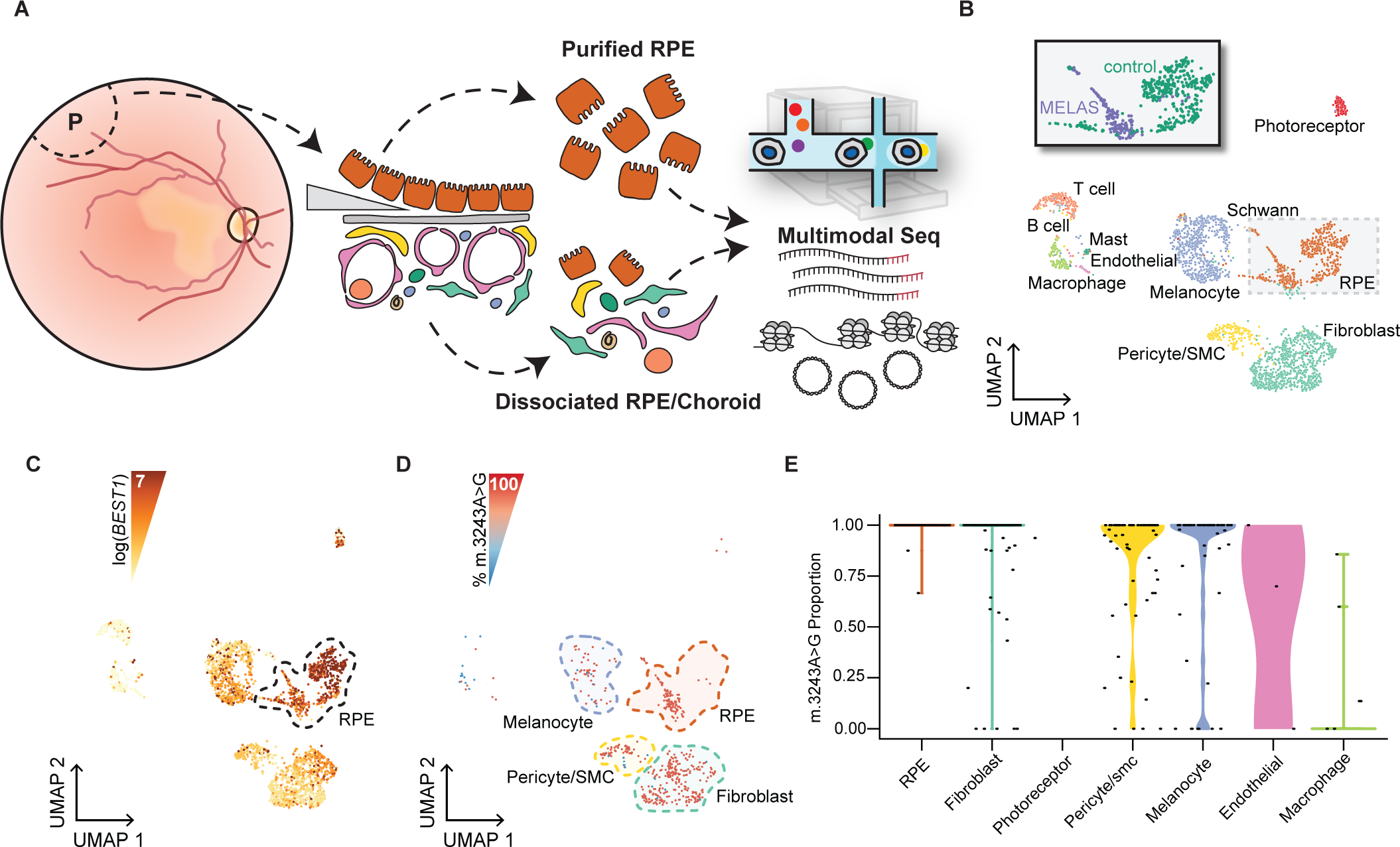
Multimodal sequencing of MELAS and control RPE/choroid. **(A)** Schematic of multimodal sequencing approach to measure gene expression, nuclear chromatin accessibility, and heteroplasmy in individual cells isolated from the RPE/choroid. **(B)** Two-dimensional UMAP embeddings of cells from the MELAS proband and two controls based on weighted nearest neighbor (WNN) analysis of scRNAseq and mt- scATACseq data. Diverging embeddings of RPE cluster between MELAS and control samples indicate abnormal gene expression (inset). **(C)** Gene expression of the RPE marker gene *BEST1*. Cells are colored based on magnitude of *BEST1* mRNA expression. **(D)** Two-dimensional UMAP embeddings of cells recovered from the proband samples. Individual cells are colored based on proportion of m.3243A>G mutant allele (red indicating 100% mutant allele and blue indicating 100% wildtype allele). **(E)** Violin plot demonstrating the proportion of m.3243A>G mutant allele in individual cells isolated from the RPE/choroid. RPE cells are homoplasmic for the mutant m.3243G allele.

We performed H&E staining on two unrelated MELAS retinal samples. Both donor sections exhibited signs of RPE dysfunction, specifically an epithelial to mesenchymal transition (EMT) that has been previously recognized in models of MELAS *in vitro* (47). MELAS donors exhibited RPE thinning (**Figure 9A-C**), a recognized signed of RPE EMT (50) as well as cells with swollen cytoplasm. Additionally, when differential gene expression between MELAS and control RPE cells were analyzed, several genes known to be RPE-specific (*BEST1*, *RLBP1*) and involved in lysosomal function (*ATP10B*) were found to be dysregulated (**Figure 9D**).

**Figure 9.**
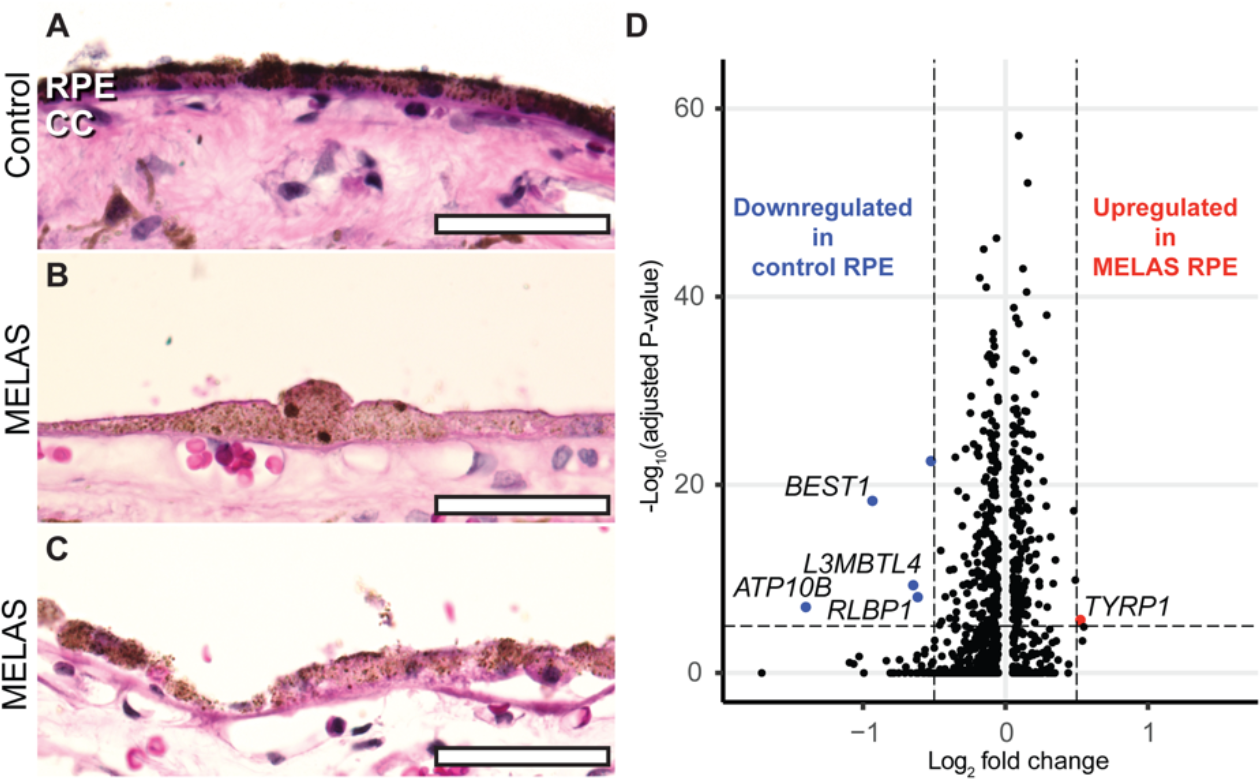
m.3243A>G-induced dysfunction in the RPE. **(A)** Hematoxylin and eosin staining of the RPE and choroid (CC) of a healthy control donor. RPE monolayer shows typical cuboidal epithelial morphology. (**B)** RPE dysplasia is observed in the proband, including both thinning that is associated with epithelial to mesenchymal transition and hypertrophy. **(C)** RPE morphology is also abnormal in an unrelated MELAS patient with the m.3243A>G variant including RPE thinning and migration into the retina. **(D)** Differential gene expression analysis of multimodal sequencing data between MELAS and control RPE cells shows several differentially expressed genes known to be involved in normal RPE function. Scale bar represents 50µm.

## DISCUSSION

In this study, we sought to probe the distribution and consequences of m.3243A>G heteroplasmy in a clinically-relevant human tissue. We profiled heteroplasmy, gene expression, and chromatin accessibility in single cells and found that m.3243G is non-randomly distributed, being high in neural retina and RPE and lower in the choroidal endothelium. We also found that the presence of m.3243G results in dysregulation of genes involved in oxidative stress management and the mTOR/PI3K/AKT signaling pathway. Differential gene expression was observed in cell types lacking pathogenic allele, implying that this pathway may have a role in heteroplasmy modulation *in vivo*. We further showed that transcription factor binding motif enrichment is dysregulated in MELAS in a cell type-specific manner, and that factors involved in mTOR signaling are differentially active in the disease state. Finally, by combining previous ophthalmologic clinical observations (24, 51) and multimodal sequencing of the RPE, we showed that the RPE dysfunction that is associated with high proportions of m.3243G is likely an early step in the pathogenic sequence that results in retinal dystrophy of MELAS and related mitochondrial conditions (i.e., MIDD and Leigh Syndrome).

### m.3243G is non-randomly distributed between ocular cell types

Mitochondria harboring pathogenic variants have long been known to be subject to purifying selection during oogenesis and early embryonic development (52). More recently, evidence has emerged that such selection may occur within certain tissues on a cell type-specific basis (10, 11, 53). Currently, the mechanism that leads to uneven mitochondrial DNA partitioning between cell types of the same individual is not clear. One hypothesis is that mitochondrial genomes are randomly partitioned into early embryonic progenitors during the first zygotic divisions. In such a model, high proportions of a mutant mtDNA in the central nervous system would arise from mutant mtDNA genomes being randomly partitioned into a neuroectoderm-fated cell in early embryonic development. However, reproducible patterns of mtDNA distribution across specific tissues of unrelated individuals (30) and within specific cell types within tissues of unrelated individuals (11) suggest that there is cell type-level control over mtDNA partitioning. Our data show that within the microenvironment of the retina and choroid, certain cell types develop lower levels of m.3243G despite a background of globally elevated m.3243G in a severely affected patient. We find that choroidal T cells possess low levels of m.3243G, building on previous observations of lower m.3243G in circulating T cells (11) and observe a similar phenomenon in the endothelial cells of the choroid.

### The m.3243A>G variant causes macular retinopathy with evidence of RPE/choroid dysfunction

It has been hypothesized that the unique macular specificity in m.3243A>G disease is caused by region-specific dysfunctional RPE cells (54). A notable anatomical feature of retinal dystrophy caused by m.3243A>G is the presence of outer retinal tubulations (ORT) (24, 34, 35), structures formed by cone photoreceptor reorganization driven by Müller cell activation (48). Clinical examination of the retina of MELAS patients suggests a disease process that begins in the RPE and/or choroid, and eventually leads to outer retinal damage. Our data showing a lack of the mutant allele in the choroidal endothelium supports the theory that the RPE is the key player in the formation of m.3243A>G-associated retinal disease.

The endothelial cells of the choriocapillaris form in early development from a pool of hemanogioblasts that also give rise to primitive blood cells (55–57). The differentiation of mesenchymal progenitors that gives rise to the choriocapillaris is distinct from the angiogenic process that later connects the choroidal circulation to that of the rest of the body (55). Notably, the choroidal fibroblasts that displayed high m.3243G burden are also derived from the same mesenchymal progenitor pool, indicating that cell type-specific selection against the mutant allele as opposed to coincidental pattern following developmental lineages. The unique developmental origin of choroidal endothelium may explain the low proportion of m.3243G observed in this study as compared to a previous study of m.3243A>G in other endothelial cell populations (58).

### Retinal m.3243A>G heteroplasmy in the context of metabolism

The metabolic program of retinal cells varies greatly by cell type (59, 60). Photoreceptor cells have the highest energy needs in the retina, primarily utilizing ATP to drive ion pumps that are essential for light modulated changes in membrane potential (60). However, photoreceptors generate most of their ATP through aerobic glycolysis as opposed to the more efficient process of oxidative phosphorylation (61). While photoreceptors contain numerous mitochondria, their primary function may be to generate biosynthetic intermediates via the TCA cycle instead of ATP via the electron transport chain. Because of this, mistranslation of electron transport chain components in MELAS may negatively affect photoreceptors to a lesser than RPE.

Lactate generated by photoreceptor glycolysis is exported and utilized by glial and RPE cells as a substrate for oxidative phosphorylation in their mitochondria (59) in a reversed process reminiscent of the neuron/astrocyte metabolic linking elsewhere in the central nervous system. The reliance of the RPE on oxidative phosphorylation is necessary to support photoreceptor cell glycolysis, given their physical arrangement with respect to the choroid. Glucose arrives to the outer retina via choroidal blood flow and must be passed through the RPE layer to reach the photoreceptors (see **Figure 3A**). Because of this arrangement, RPE preference for oxidative phosphorylation using lactate and other fuels allows for minimal glucose loss during transport to the photoreceptors under normal conditions (61). Additionally, RPE energy demand is supplemented by oxidation of ingested photoreceptor outer segment fatty acids (62).

Cell type-specific metabolic pathway preference appears to underlie the observed distribution of m.3243A>G burden across the retina and choroid and the pathogenic mechanism of the variant in the retina. The m.3243A>G variant is thought to impair tRNA charging by cognate aminotransferases (15, 63) and as a result, likely disrupts electron transport chain protein translation (18). These changes in turn disrupt the process of oxidative phosphorylation in cells with high levels of m.3243A>G (20). It has been shown previously that cells harboring high levels of m.3243G are predisposed to favor glycolysis(42). Photoreceptor cells do not largely rely on the electron transport chain for energy production, and as such may be more tolerant of the accumulation of mtDNA carrying the pathogenic allele. However, a high proportion of m.3243G in the RPE, as seen in the current study, likely causes decreased electron transport chain function and leads to derangement of the RPE-photoreceptor cooperative metabolic ecosystem, eventually causing photoreceptor dysfunction and vision loss.

### m.3243A>G-induced changes transcription and chromatin accessibility involve mTOR signaling

The non-random distribution of heteroplasmy observed in the eye invites the question of how heteroplasmy may be modulated in a cell type-specific manner. Understanding of such a phenomenon would open new avenues for treatment of mitochondrial disease. Previous work has shown that chemical inhibition of the mTOR/PI3K/AKT signaling axis is sufficient to reduce the proportion of m.3243G allele in cultured cybrid cell lines(42). In our analysis of differential gene expression, we discovered that differentially expressed gene lists of cell types with both high and low m.3243G burdens were enriched for genes targeted by the same mTOR/PI3K/AKT inhibitors (i.e., sirolimus, LY294002, torin1) as well as endogenous modulators of the pathway (i.e., LARP1 & RICTOR). Motif enrichment analysis of mt-scATACseq data further showed dysregulation of transcriptional signaling downstream of mTOR in MELAS cells in low-heteroplasmy cell types in changes of YY1 and NR3C2 motif enrichment. Together, these analyses implicate mTOR signaling in the modulation of heteroplasmy *in vivo*, building on previously observations of its sufficiency to control heteroplasmy *in vitro*.

### Limitations of the current study

The current study has limitations that necessitate future experimental work. Though m.3243A>G is the most common pathogenic mtDNA variant with an estimated individual frequency of 0.018%(64), cases of visual symptoms caused by this variant are still relatively rare, with an estimated 0.0002% of the population of the United States affected(65). As such, it is exceedingly difficult to acquire appropriate biological replicates of post-mortem ocular tissues suitable for single-cell analysis. We validated our mt-scATACseq findings of non-random heteroplasmy distribution using LCM-dPCR on independent biological samples (**Figure 4**). The data in the current study is further supposed by previously observations of purifying selection against the m.3243G allele in circulating T cells. We observed the same trend in the choroidal T cells captured in this study, lending credence to our findings in other ocular cell types.

In this study, we focused only on the most common pathogenic mitochondrial variant. While our results imply a generalizable phenomenon for the establishment of mitochondrial variant segregation in the retina and choroid, further study is needed to clarify whether variants segregate in a generalized or variant-specific manner. The cell type-specificity of pathogenesis in other retinal diseases caused by mtDNA variants such as Leber Hereditary Optic Neuropathy (LHON) and Neuropathy, Ataxia, and Retinitis Pigmentosa (NARP) provides evidence that some heteroplasmy modulation may be variant-specific. Such knowledge may also clarify the underlying mechanism of mtDNA segregation within tissues.

### Conclusions

In this study, we used a single-cell approach to measure m.3243G heteroplasmy in single cells of a developmentally complex tissue. We found that m.3243G was lowest in the endothelial cells of the choriocapillaris of two MELAS patients and highest in the neural retina and RPE. We also found that in cells with high m.3243G chromatin accessibility and gene expression is altered. Together, these data support the hypothesis that high m.3243G in the RPE initiates MELAS-related retinal dystrophy. Observation of non-random m.3243G partitioning in adult samples implies selection against heteroplasmy in certain cell types during development. Ultimately, elucidation of how certain cells can regulate m.3243A>G heteroplasmy and other variants may lead to new therapeutic approaches for MELAS and other mitochondrial diseases.

## METHODS

### Ocular tissue acquisition

Human donor eyes used in single cell studies from a patient affected with MELAS and an unaffected donor were acquired from the Iowa Lions Eye Bank. A second pair of eyes from a donor affected with MELAS was obtained from the Minnesota Lions Eye Bank. All eyes used in the study were obtained or preserved within 9 hours of death and all experiments were performed in compliance with the Declaration of Helsinki and following full consent of the donor’s next of kin. Retinal and choroidal tissues were isolated from 8-mm trephine punch biopsies of the macular and inferotemporal peripheral regions and processed as previously described (38).

Briefly, the neural retina was dissected away from the underlying retinal pigment epithelium and choroid prior to tissue dissociation. Neural retina samples were dissociated with 20 units/mL of papain (Worthington Biochemical Corporation) with 0.005% DNase I (Worthington Biochemical Corporation) on a shaker at 37°C for 75 minutes. RPE/choroid samples were dissociated mechanically using razor blades into one-millimeter pieces then dissociated enzymatically using collagenase on a shaker at 37°C for 60 minutes. Following dissociation, cells were cryopreserved in DMSO-based Recovery Cell Cryopreservation Media (Gibco). After overnight cooling at -80°C, samples were transferred to liquid nitrogen (vapor phase) for long-term storage.

### Single-cell gene expression (scRNAseq) library construction

Cryopreserved cells were rapidly thawed at 37°C and resuspended in dPBS-/- (Gibco) with 0.04% non-acetylated bovine serum albumin (New England Biolabs). Cells were filtered through a 70µm filter and diluted to target 8,000 cells per run. Single cells were then partitioned and barcoded with the Chromium Controller instrument (10X Genomics) and Single Cell 3’ Reagent (v3.1 chemistry) kit (10X Genomics) according to the manufacturer’s specifications with no modification (Rev C). Final libraries were quantified using the Qubit dsDNA HS Assay Kit (Life Technologies) and diluted to 3ng/µL in buffer EB (Qiagen). Library quality was confirmed using the Bioanalyzer High Sensitivity DNA Assay (Agilent) prior to sequencing.

### Mitochondrial single-cell ATAC (mt-scATACseq) library construction

Cells were prepared for mt-scATAC sequencing using the protocol described previously by Lareau et al. (37). Briefly, cells were thawed as above and filtered through a 70µm cell strainer. Cells were then washed twice with dPBS-/- (Gibco). Cells were fixed with 1% formaldehyde (Sigma) in PBS-/- for 10 minutes at room temperature. Fixation was quenched with the addition of glycine (Research Products International) to a final concentration of 0.125M for 5 minutes at room temperature. Following fixation, cells were washed twice with ice-cold PBS-/-. Following the second wash, cells were resuspended in 100µL ice-cold Lysis Buffer (10 mM Tris-HCl pH 7.4 (Millipore-Sigma), 10 mM NaCl (Ambion/Invitrogen), 3 mM MgCl2 (Ambion/Invitrogen), 0.1% Nonidet-P40 substitute (Research Products International), 1% Fraction V BSA (Research Products International)) and incubated on ice for 2.5 minutes. Following incubation, 1mL of ice-cold Wash Buffer (10 mM Tris-HCl pH 7.4, 10 mM NaCl, 3 mM MgCl2, 1% BSA) was added, and cells were pelleted at 500 x g for 5 minutes at 4°C. Cells were resuspended in 10µL diluted Nuclei Buffer (10X Genomics) and counted using Trypan Blue (Gibco) with the Countess automated cell counter (Invitrogen). Based on these counts, cell suspensions were diluted in diluted Nuclei Buffer to a final target concentration of 6,000 cells per µL. Cells were then processed following the 10X Chromium Next GEM Single Cell ATAC Reagent Kit v1.1 User Guide (Rev. D) without modification. Final libraries were quantified using the Qubit dsDNA HS Assay Kit (Life Technologies) and diluted to 3ng/µL. Library quality was checked using the Bioanalyzer instrument (Agilent), wherein periodicity of fragment length was observed, consistent with high quality ATAC-seq library construction.

### Multimodal mt-scATAC and scRNA library preparation

Cryopreserved RPE & choroid samples were thawed as described above. Cells were prepared for multimodal profiling using the sample preparation modifications described previously (66). Briefly, cells were thawed and filtered through a 70µm cell strainer. Cells were then washed twice with dPBS-/- (Gibco). Cells were fixed with 1% formaldehyde (Sigma) in PBS-/- for 10 minutes at room temperature. Fixation was quenched with the addition of glycine (Research Products International) to a final concentration of 0.125M for 5 minutes at room temperature. Following fixation, cells were washed twice with ice-cold PBS-/-. Following the second wash, cells were resuspended in 100µL ice-cold Lysis Buffer (10 mM Tris-HCl pH 7.4 (Millipore-Sigma), 10 mM NaCl (Ambion/Invitrogen), 3 mM MgCl2 (Ambion/Invitrogen), 0.1% Nonidet-P40 substitute (Research Products International), 1% Fraction V BSA (Research Products International)) and incubated on ice for 2.5 minutes.

Following incubation, 1mL of ice-cold Wash Buffer (10 mM Tris-HCl pH 7.4, 10 mM NaCl, 3 mM MgCl2, 1% BSA) was added, and cells were pelleted at 500 x g for 5 minutes at 4°C. Cells were resuspended in 10µL diluted Nuclei Buffer (10X Genomics) and counted using Trypan Blue (Gibco) with the Countess automated cell counter (Invitrogen). Cells were then processing following the 10X Genomics Chromium Next GEM Single Cell Multiome ATAC + Gene Expression (Rev. E) without modification. Final libraries were quantified using the Qubit dsDNA HS Assay Kit and diluted to 3ng/µL. Library quality was checked using the Bioanalyzer system.

### scRNA-seq sequencing, preprocessing, and mapping

scRNA libraries were pooled and sequenced using the NovaSeq 6000 instrument (Illumina) generating 100-bp paired end reads. Sequencing was performed by the Genomics Division of the Iowa Institute of Human Genetics. FASTQ files were generated from base calls with the bcl2fastq software (Illumina), and reads were mapped to the pre-built GRCh38 reference with Cell Ranger v3.0.1 (10X Genomics) using the ‘count’ function. Only cells passing the default Cell Ranger call were analyzed further. Neural retina and RPE/choroid samples were integrated separately using canonical correlation analysis (CCA) in Seurat v3.1 (40). Only cells with between 500 and 7,000 unique genes (features) were included in the analysis. For RPE/choroid samples, only cells with < 65% of reads mapping to mtDNA-encoded genes and < 40% of reading mapping to ribosomal RNA genes were included. For neural retina samples, only cells with < 40% of reads mapping to mtDNA-encoded genes and < 20% of reading mapping to ribosomal RNA genes were included.

### mtscATAC-seq sequencing, preprocessing, and mapping

mt-ATAC libraries were pooled and sequenced using the Illumina NovaSeq 6000 instrument generating 100-bp paired end reads. Sequencing was carried out by the Genomics Division of the Iowa Institute of Human Genetics. FASTQ files were generated from base calls with the bcl2fastq software (Illumina). mtscATAC-seq reads were mapped to a modified GRCh38 reference genome that was generated by hard-masking nuclear regions that share homology with mitochondrial sequences, as described previously (37). Mapping and quantification were carried out by Cell Ranger ATAC (v2.0.0, 10X Genomics) using the ‘count’ function, while the mt-ATAC portion of the multimodal library reads were mapped using Cell Ranger ARC (v.2.0, 10X Genomics). Only cells passing the default Cell Ranger call were analyzed further. Single-cell heteroplasmy of mitochondrial variants was determined using the mgatk package (v0.6.1).

Neural retina and RPE/choroid samples were integrated separately using Signac v1.4.0 (40). Briefly, integration anchors were identified with reciprocal LSI projection. LSI embeddings were then integrated using the ‘IntegrateEmbeddings’ function, and a UMAP was generated. Only cells with peak region fragments between 1,000 and 75,000 and average mtDNA depth between 10x and 1,000x were used for integration and in downstream analyses. Manually annotated cluster labels were transferred from the scRNA-seq data to the integrated mt-scATAC-seq data. First, transfer anchors between the integrated scRNA-seq and mt-scATAC-seq objects were identified using the ‘FindTransferAnchors’ function. Next, predicted cluster labels were generated using the TransferData function. Only cells annotated with a prediction.score > 0.6 were used in subsequent analysis.

### Multimodal sequencing, preprocessing, and mapping

FASTQ files were generated from base calls with the bcl2fastq software (Illumina). Sequencing reads were mapped to a modified GRCh38 reference genome that was generated by hard-masking nuclear regions that share homology with mitochondrial sequences, as described previously (37). Mapping and quantification were carried out by Cell Ranger ARC (v2.0.0, 10X Genomics) using the ‘count’ function.

For gene expression libraries, cells were filtered with the Seurat (v3.1) subset function. Cells with nUMIs less than 100 (to remove cells with poor read quality) or greater than 4000 (to remove cells likely to be doublets)were removed. RPE/choroid cells with greater than 40% of reads originating from mitochondrial genes were also removed. Cells with greater than 40% of reads originating from ribosomal RNA genes were also removed. For each of the gene expression libraries, reads were normalized with the Seurat (v3.1) NormalizeData function, and variable features were found with the FindVariableFeatures command. Integration anchors from the first 25 dimensions of the canonical correlation analysis (CCA) were used to integrate gene expression data. Data scaling, principal component analysis, and clustering of the gene expression data were performed using Seurat (v3.1) (40).

For ATAC libraries, cells were filtered with the Signac (v1.4.0) subset function. Cells with fewer than 1000 peak region fragments or greater than 30000 peak region fragments were removed. Cells with greater than 1000x mtDNA depth were also removed. Mitochondrial DNA sequencing depth (mtDNA_depth) was calculated using mgatk. Term frequency inverse document frequency (TF-IDF) normalization and partial singular value decomposition were performed on each ATAC library using RunTFIDF() and RunSVD() (Signac) with default settings. Gene activities were computed using GeneActivity().

Integration anchors between ATAC datasets were identified using reciprocal LSI projection using FindIntegrationAnchors() in Signac. Dimensions 2-30 from the CCA were used. LSI embeddings were integrated using IntegrateEmbeddings based on these anchors and using dimensions 1-30. A UMAP was generated using the integrated embeddings. Cell identity within the gene expression portion of the experiment was inferred using the TransferData() function in Seurat from the annotated RPE/choroid scRNAseq data obtained in this study (Figure 2C, E). Transfer anchors between were identified using FindTransferAnchors with the following parameters: reduction = ’pca’, dims = 1:30. Cells were annotated with a prediction score > 0.8.

Gene expression and ATAC portions of the experiment were combined using Weighted Nearest Neighbor (WNN) analysis (Seurat). Integrated ATAC and gene expression objects were subset to include only common cells (i.e., cells that passed both ATAC and gene expression quality control thresholds). UMAP and cluster identification were performed on the resulting object. Cluster identification used the SLM algorithm with a resolution of 1 (Seurat). Following multimodal integration, cluster identities (i.e., cell type annotation) were refined manually based on canonical marker gene expression.

### Single cell heteroplasmy analysis

Per-cell m.3243A>G heteroplasmy was determined as described by Walker et al. (11). mt-scATACseq data were subset to retain only cells with coverage of the m.3243 locus greater than 10x and less than 1.5 times the upper boundary of the interquartile range. Multimodal sequencing data were subset to retain only cells with coverage of the m.3243 locus greater than 5x and less than 1.5 times the upper boundary of the interquartile range. Only the proband (i.e., not control) patient was included in m.3243A>G heteroplasmy analysis. Of these subsets, m.3243A>G proportion was plotted in violin plots by cell type or as a color gradient on UMAP plots.

### Differential gene expression analysis

Differential gene expression based on scRNAseq data was calculated between MELAS and control samples for each cell type annotated. The ‘FindMarkers()’ function (Seurat v4.0.3) was used with default parameters. The difference in the percent of cells expressing a given gene (‘Δ Percent Expressed’) was calculated for each gene in each cell type. Genes labeled in **Figure 6** had a log(Fold Change) ≥ 1 and Δ Percent Expressed’ ≥ 0.25. Differential gene expression based on the scRNAseq portion of the multimodal sequencing data was performed

### Pathway enrichment analysis

All genes with an adjusted p value ≤ 0.05 from the above differential expression analysis were used for pathway enrichment analysis with Ingenuity Pathway Analysis (version 01-21-03, Qiagen). Enriched canonical pathways were identified and upstream analysis was performed to identify regulators known to activate gene expression patterns similar to those observed in enriched genes. All regulators with an activation z-score ≥ 2 were plotted in **Figure 6**.

### Transcription factor binding motif enrichment analysis

Transcription factor binding motif enrichment (“ChromVAR Motif Activity”) was computed on a per-cell basis using ChromVAR run in Signac (44). The JASPAR 2020 database of motif position frequencies was used (67). Differential motif enrichment analysis was performed using the FindMarkers function of Seurat (4.0.5), using a logistic regression model and likelihood ratio test (test.use = ’LR’) with the number of ATAC counts as latent variables (latent.vars = ’nCount_ATAC’). Motif enrichment volcano plots were generated using EnhancedVolcano (1.8.0). Motif footprint analysis was carried out using the Footprint function of Seurat using the UCSC hg38 reference genome and default parameters.

### Transmission electron microscopy

Donor samples were fixed in ½ strength Karnovsky’s fixative and processed for TEM as described previously (34). Sections were cut at 85 nm with an ultra-microtome (Leica) and collected on copper slot grids coated with Formvar (0.5%) and images collected on a Hitachi HT7800 transmission electron microscope.

### Laser Capture Microdissection

Fixed frozen blocks containing neural retina and choroid from both MELAS donors and an unaffected control donor were sectioned at 7µm thickness onto membrane slides (Zeiss). Prior to sectioning, slides were exposed to UV light for 30 minutes in a tissue culture hood, per manufacturer’s suggestion. Mounted sections were rinsed once with cold deionized water prior to laser capture microdissection (LCM). LCM was carried out using the PALM Combi LCM system (Zeiss). Regions of interest (i.e., photoreceptors, RPE, or choriocapillaris) were traced and captured into adhesive cap tubes following the manufacturer’s recommended protocol. DNA was isolated using the Arcturus PicoPure DNA Extraction Kit (Applied Biosystems). Briefly, samples were digested overnight in a Proteinase K solution at 65°C, followed by a 10-minute inactivation at 95°C. DNA was stored undiluted at -20°C.

### Genotyping by digital PCR

Custom probe-based assays were designed against either variant at the m.3243 locus as previously described (30). Probes were designed incorporating several locked nucleic acids (indicated with ‘+’ below) to attain adequate binding specificity to provide allelic specificity (**Table 3**) and manufactured by Integrated DNA Technologies (Coralville, IA). Digital PCR was carried out using the 3D PCR platform (Applied Biosystems).

For the m.3243A/G assay, the following mixture was used (per reaction): 8.7µL 3D Master Mix V2 (Applied Biosystems), 0.85µL A-FAM Probe (5µM), 0.85µL G-HEX Probe (5µM), 1.67µL F Primer (5µM), 1.67 R Primer

(5µM), 1.66µL H2O, 1µL DNA lysate from LCM. 14.5µL of the mixture was loaded into a 3D Digital PCR chip using the QuantStudio Digital PCR Chip Loaded following the manufacturer’s instruction.

The thermal cycler program was run as follows for the m.3243A/G assay: Step 1: 95°C – 10 minutes; Step 2: 56°C – 2 minutes; Step 3: 98°C – 30 seconds; Step 4: 60°C – 2 minutes, with steps 2 and 3 cycled 45 times. After amplification, chips were allowed to equilibrate to room temperature for approximately 10 minutes and were then read. Data were analyzed using the 3D Quantstudio cloud-based web application, and the automatically generated thresholds were used. The proportion of G allele for the *MT-TL1* A/G heteroplasmy assay was calculated as:

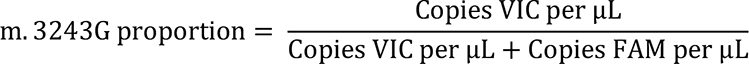

### Isolation of Primary Retina Pigmented Epithelium (RPE)

Donor eyes were dissected as previously described (39). Following removal of the vitreous and neural retina, the RPE was manually scraped away from the underlying Bruch’s membrane and choroid. Cells were transferred directly to cryopreservation media and cryopreserved in the same manner as described for the dissociated neural retina and RPE/choroid above.

### Human donor samples

Human donor sample characteristics are summarized in **Table 2**.

**Table 2.** Donor characteristics and experimental details.

| Donor ID | Age | Sex | PMI (h:m) | Experiment | Genotype | Region |
| --- | --- | --- | --- | --- | --- | --- |
| MELAS 1 | 41 | Male | 8:09 | 1, 2, 3, 4, 5, 7 | m.3243A>G | Macula + Periphery |
| MELAS 2 | 70 | Female | 7:38 | 5, 6, 7 | m.3243A>G | Macula |
| Control 1 | 40 | Male | 5:40 | 1, 2 | n/a | Macula + Periphery |
| Control 2 | 59 | Male | 6:59 | 3 | n/a | Periphery |
| Control 3 | 60 | Male | 6:14 | 4 | n/a | Periphery |
| Control 4 | 70 | Female | 5:22 | 5 | n/a | Macula |
| Control 5 | 83 | Male | 8:30 | 6 | n/a | Macula |
| Control 6 | 77 | Male | 8:00 | 6 | n/a | Macula |
| Control 7 | 41 | Female | 7:38 | 7 | n/a | Macula |

1. scRNAseq 2. mt-scATACseq 3. Multimodal profiling (RPE/Choroid) 4. Multimodal profiling (Purified RPE) 5. LCM-dPCR 6. Transmission electron microscopy 7. Hematoxylin and eosin staining

**Table 3.**
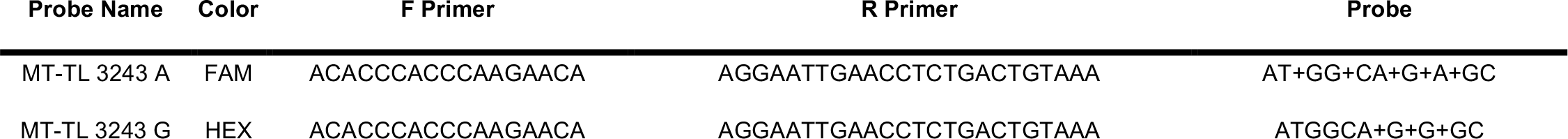
Digital PCR assay details.

### Statistics

Differentially expressed genes between disease conditions were identified using a Wilcoxon Rank Sum test. The p value was adjusted based on Bonferroni correction using all genes in the dataset. Differential motif enrichment analysis was performed using a logistic regression model and likelihood ratio test with the number of ATAC counts used as latent variables.

### Study approval

Human donor eyes used in this study were acquired from the Iowa Lions Eye Bank or the Minnesota Lions Eye Bank in accordance with the Declaration of Helsinki following consent of the donor’s next of kin. The human subjects aspects of the study were approved by the Institutional Review Board of the University of Iowa. The patients for whom clinical data is available were seen at the University of Iowa Hospitals and Clinics and provided written informed consent for inclusion in this study.

## DATA ACCESS

All raw and processed sequencing data generated in this study have been submitted to the NCBI Gene Expression Omnibus (GEO; https://www.ncbi.nlm.nih.gov/geo/) under accession number GSE202747.

## AUTHOR CONTRIBUTIONS

N.K.M., A.P.V., B.A.T., and R.F.M. designed the study. N.K.M., A.P.V., M.J.F.-W., X.L., and K.V. conducted experiments and acquired data. E.M.S. examined the proband and provided clinical data. N.K.M. and A.P.V. analyzed the data. N.K.M. wrote the original draft of the manuscript. A.P.V., E.M.S., B.A.T., and R.F.M. edited the manuscript. All authors reviewed and approved the final manuscript.

## ACKNOWLEDGEMENTS

This work was supported by the National Institutes of Health (T32GM008629, T32GM139776, F30EY034009, P30EY025580, F30EY031923, R01EY033308), Research to Prevent Blindness, the Elmer and Sylvia Sramek Charitable Trust, and The Edward N. & Della L. Thome Memorial Foundation. The authors would like to acknowledge the extraordinary generosity of the tissue donors and their families that made this study possible. We would also like to thank K. Mulfaul, E. Burnight and the other members of the Tucker and Mullins laboratories for their helpful conversation and advice. Data presented herein were obtained at the Genomics Division of the Iowa Institute of Human Genetics which is supported, in part, by the University of Iowa Carver College of Medicine. We would like to acknowledge specifically the advice and support of M. Boes, G. Hauser, K. Knudtson, A. Sheehan, and E. Snir. Colors used in plots were based on www.colorbrewer2.org by Cynthia A. Brewer, Department of Geography, Pennsylvania State University.

