## Supplemental Figures for "Multimodal single-cell analysis of non-random heteroplasmy distribution in human retinal mitochondrial disease"

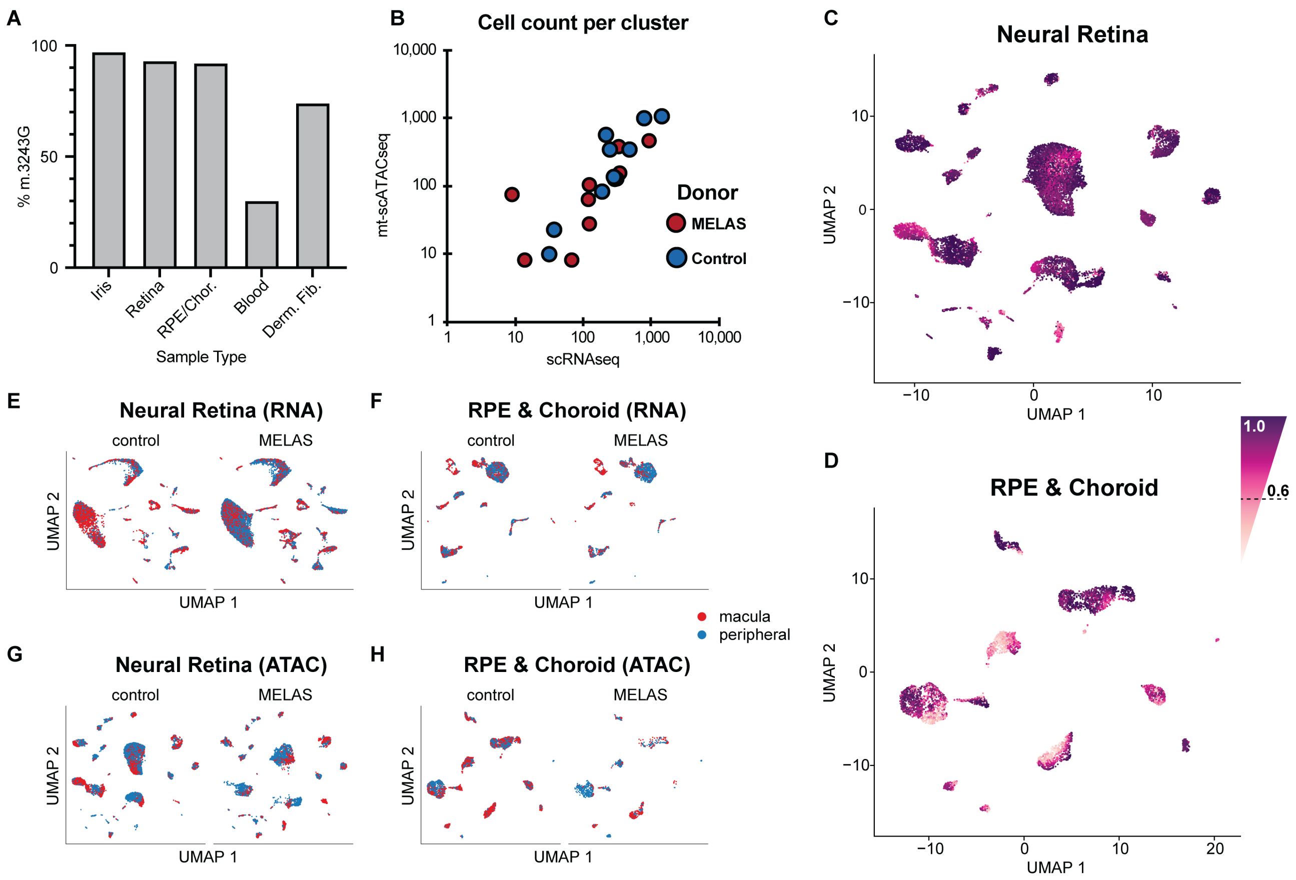


**Figure S1. scRNAseq and mt-scATACseq of MELAS and control ocular cells. (A)** m.3243A>G heteroplasmy as measured by digital PCR in ocular and non-ocular cell types from the proband (MELAS 1). **(B)** High correlation in cell number per cluster as measured by scRNAseq and mt-scATACseq modalities across MELAS and control samples. Each dot represents one cell type in the conrol (blue) or MELAS (red) donor. **(C)** Two-dimensional UMAP embedding of neural retinal cells based on mt-scATACseq data from the proband and control donor. Cells are colored based on label transfer prediction score. **(D)** Two-dimensional UMAP embedding of RPE and choroidal cells based on mt-scATACseq data from the proband and control donor. Cells are colored based on label transfer prediction score. **(E, F)** Two-dimensional UMAP embedding of neural retinal and RPE/choroidal cells based on scRNAseq data. **(G, H)** Two-dimensional UMAP embedding of neural retinal and RPE/choroidal cells based on mt-scATACseq data. Cells in **(E-H)** are split by donor and colored based on region of sample.

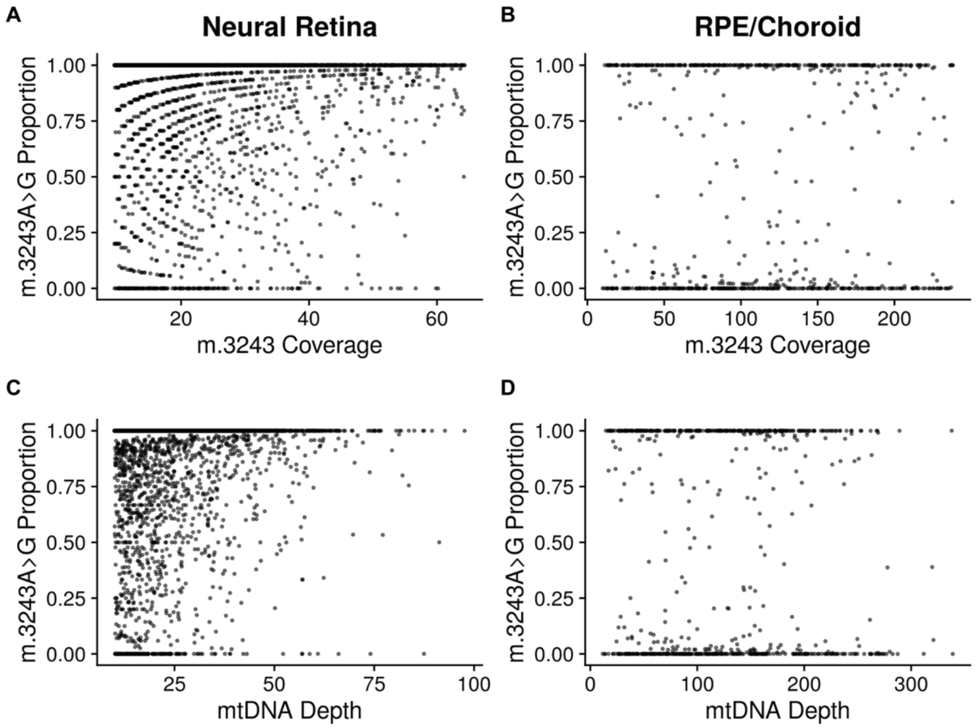


**Figure S2. m.3243A>G heteroplasmy measurement by mt-scATACseq in the context of locus and genome coverage. (A, B)** Each cell from **Figure 3** is plotted with measured m.3243G proportion on the y-axis and mt-scATACseq coverage of the m.3243 locus on the x-axis. No correlation is observed; cells with both high and lower coverage of the m.3243G locus display a range of m.3243G proportions. **(C, D)** Cells from **Figure 3** are shown with m.3243G proportion plotted against average mtDNA sequencing depth. As in **A** and **B**, no correlation is observed, showing that neither sequencing depth nor m.3243 coverage by mt-scATACseq confounds measurement of m.3243A>G heteroplasmy.

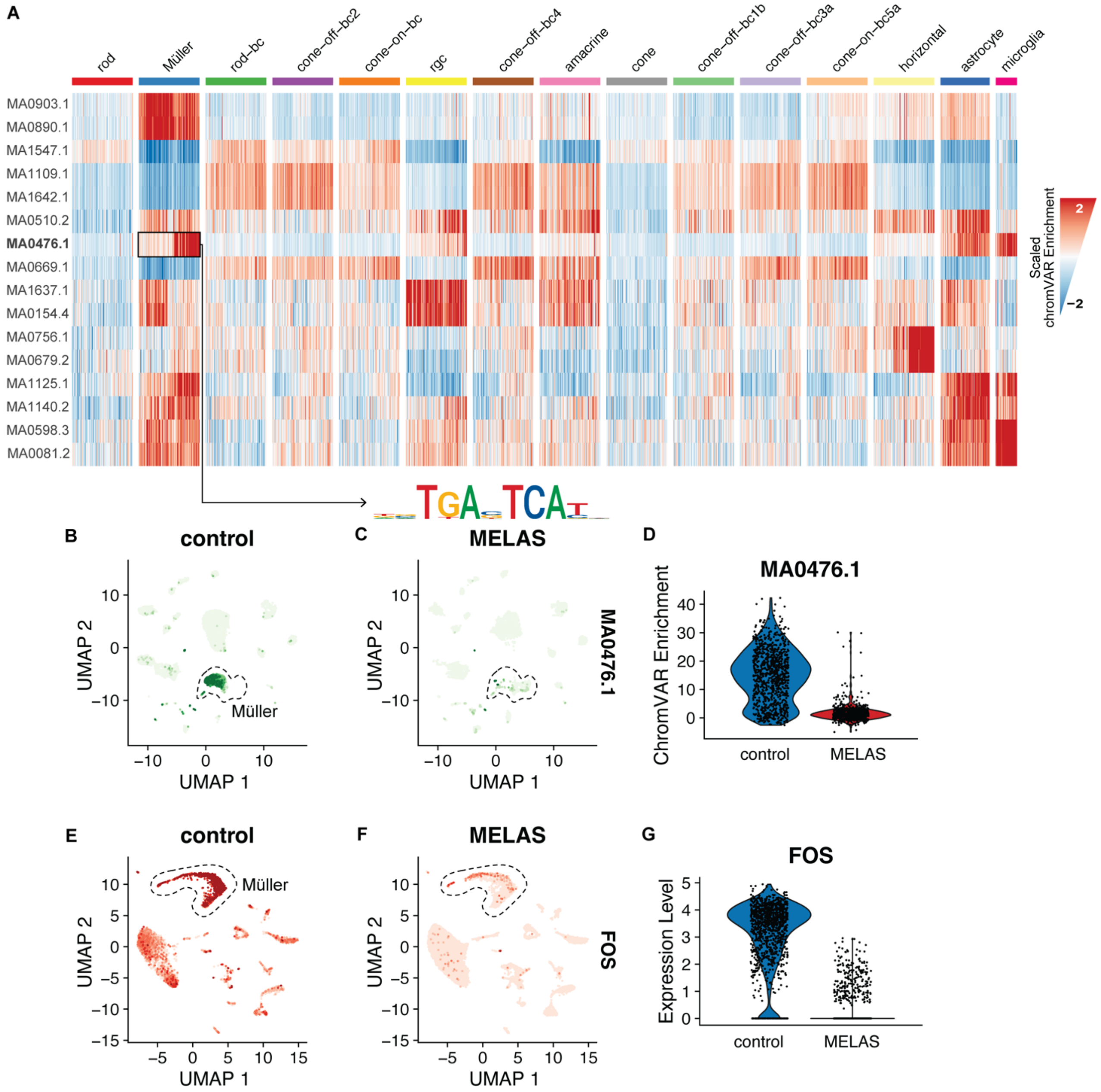


**Figure S3. Cell type and disease state-specific transcription factor binding motif enrichment. (A)** Heatmap of scaled chromVAR enrichment scores for top cell type-specific motifs. A sample of no more than 100 cells are shown for each cell type. JASPAR matrix names corresponding to marker motifs are shown. **(B, C)** UMAP of neural retinal cells colored by chromVAR enrichment score for the MA0476.1/FOS motif. Enrichment is shown in the control Müller cell cluster but not MELAS. **(D)** Violin plot of MA0476.1/FOS motif enrichment in the Müller cell cluster. **(E, F)** scRNA-seq drive UMAP of neural retinal cells colored by *FOS* transcript abundance. Downregulation is shown in the MELAS Müller cell cluster compared to control. **(G)** Downregulation of *FOS* expression in MELAS Müller cells.

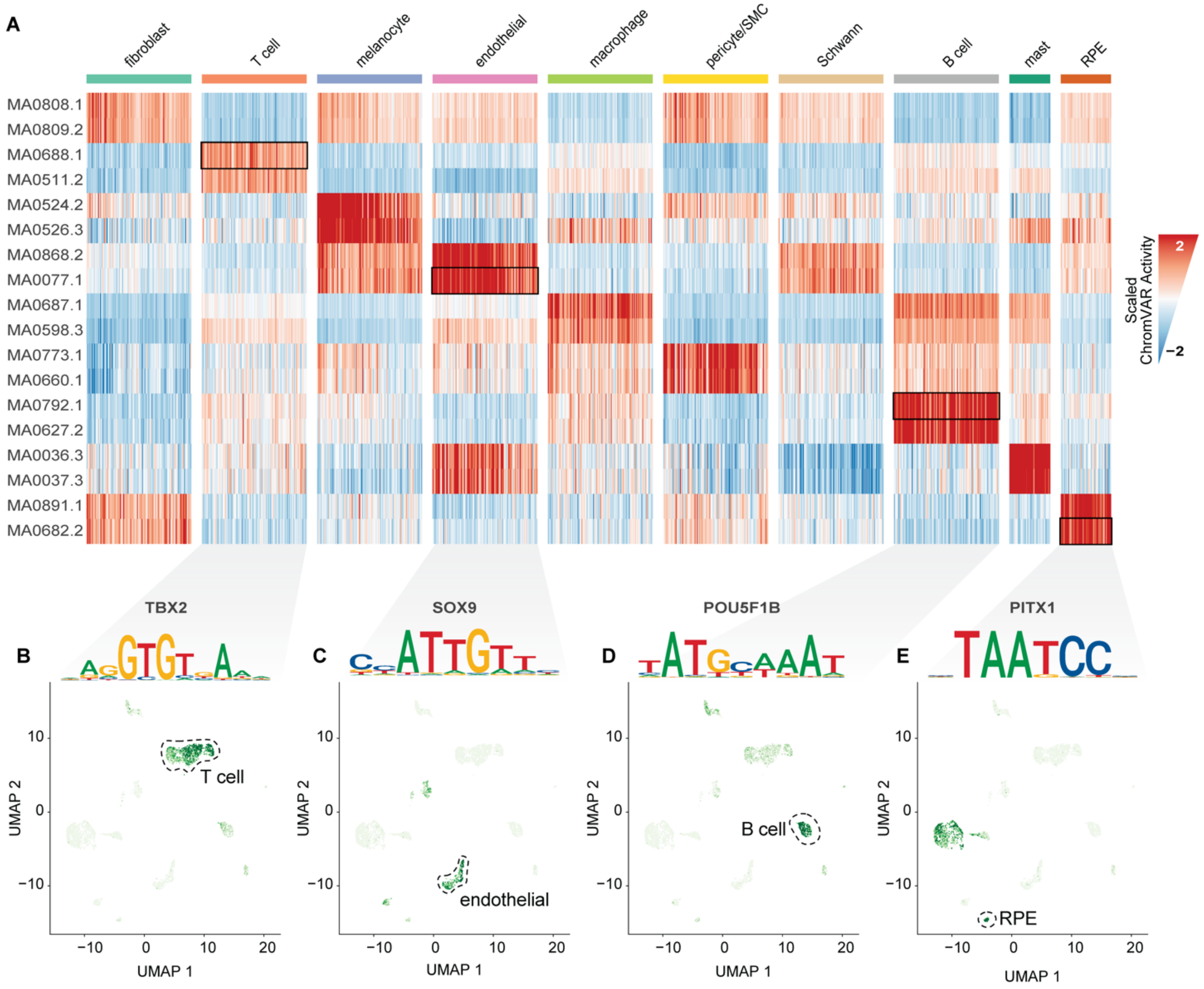


**Figure S4. Cell type defining motif enrichment in the choroid. (A)** Heatmap displaying the chromVAR enrichment score of marker JASPAR motifs for the nine choroidal and RPE cell types captured by mt-scATACseq. **(B)** *TBX2* binding motif is enriched in T cells as shown by chromVAR score overlaid onto single cells in the UMAP. **(C)** *SOX9* binding motif is enriched in endothelial cells. **(D)** *POU5F1B* binding motif is enriched in B cells. **(E)** *PITX1* binding motif is enriched in RPE.

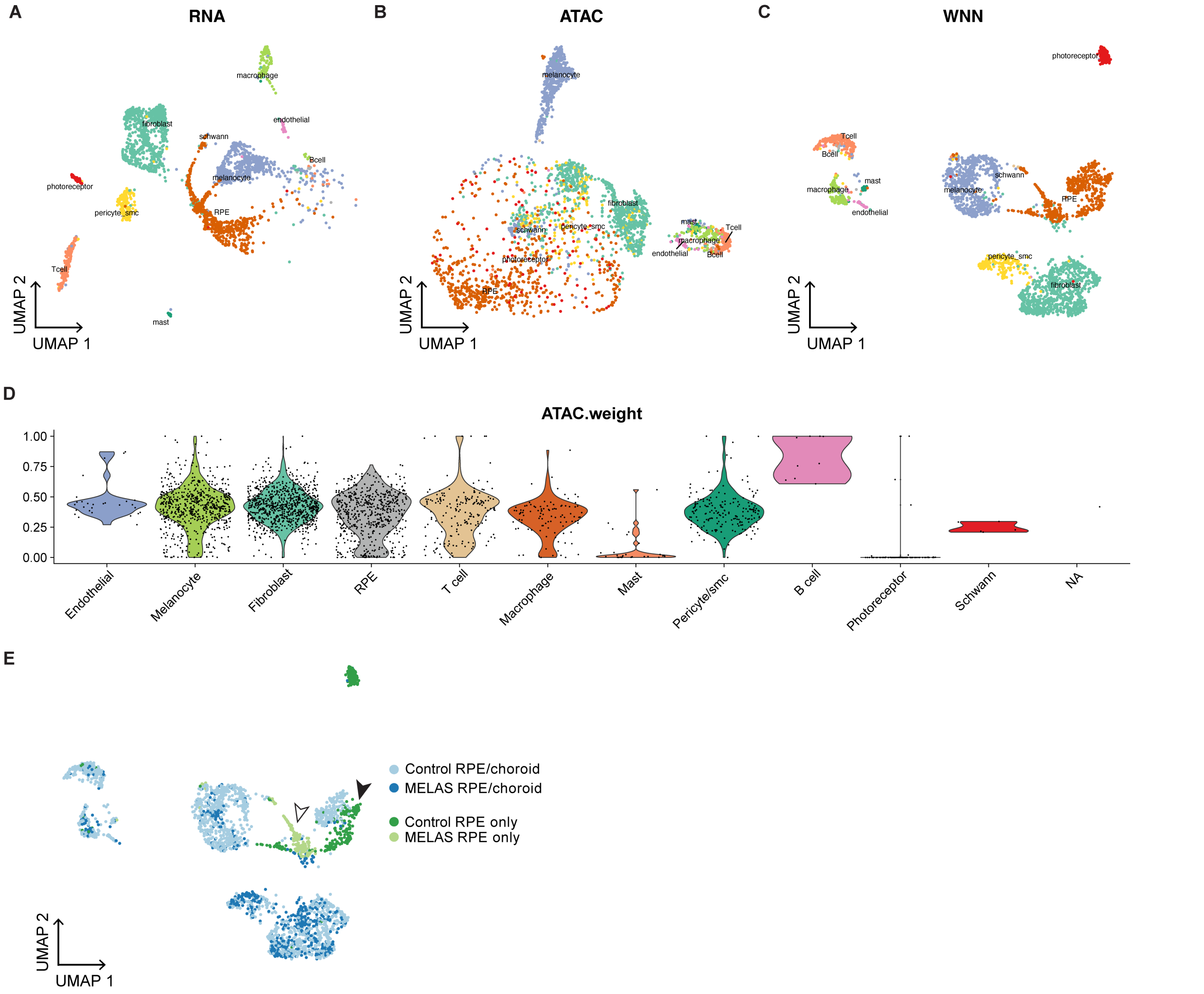


**Figure S4. Integration of multimodal data from choroidal and RPE single cells.** Weighted nearest neighbor analysis was used to combine sequencing modalities and drive dimensionality reduction and clustering. **(A)** Two-dimensional UMAP embedding of RPE/choroid cells based on scRNAseq modality. **(B)** Two-dimensional UMAP embedding of RPE/choroid cells based on mt-scATACseq modality. **(C)** Two-dimensional UMAP embedding of RPE/choroid cells based on weighted scRNAseq and mt-scATACseq modalities. **(D)** Modality weighting in (C) per cell, shown grouped by cell type. **(E)** Two-dimensional UMAP embedding of RPE/choroid cells based WNN analysis. Cells are colored based on donor disease state (MELAS or control) and sample category (whole RPE/choroid or purified RPE). In general, clustering between disease state and sample category is observed. In the RPE cluster, the MELAS (white arrowhead) and control (black arrowhead) RPE samples segregate away from each other indicating diverging states.
